# Exploring ensemble structures of Alzheimer’s amyloid β (1-42) monomer using linear regression for the MD simulation and NMR chemical shift

**DOI:** 10.1101/2021.08.23.457317

**Authors:** Wonjin Yang, Beom Soo Kim, Yuxi Lin, Dai Ito, Jin Hae Kim, Young-Ho Lee, Wookyung Yu

**Affiliations:** Department of Brain and Cognitive Sciences, DGIST, 333 Techno jungang-daero, Daegu 42988, Republic of Korea; Research Center for Bioconvergence Analysis, Korea Basic Science Institute, Ochang, Chung Buk 28119, Republic of Korea; Department of New Biology, DGIST, 333 Techno jungang-daero, Daegu 42988, Republic of Korea; Department of Bio-analytical Science, University of Science and Technology, Daejeon 34113, Republic of Korea; Graduate School of Analytical Science and Technology, Chungnam National University, Daejeon 34134, Republic of Korea; Research headquarters, Korea Brain Research Institute, Daegu 41068, Republic of Korea; Core Protein Resources Center, DGIST, 333 Techno jungang-daero, Daegu 42988, Republic of Korea

## Abstract

Aggregation of intrinsically disordered amyloid β (Aβ) is a hallmark of Alzheimer’s disease. Although complex aggregation mechanisms have been increasingly revealed, structural ensembles of Aβ monomers with heterogeneous and transient properties still hamper detailed experimental accesses to early events of amyloidogenesis. We herein developed a new mathematical tool based on multiple linear regression to obtain the reasonable ensemble structures of Aβ monomer by using the solution nuclear magnetic resonance (NMR) and molecular dynamics simulation data. Our approach provided the best-fit ensemble to two-dimensional NMR chemical shifts, also consistent with circular dichroism and dynamic light scattering analyses. The major monomeric structures of Aβ including β-sheets in both terminal and central hydrophobic core regions and the minor partially-helical structures suggested initial structure-based explanation on possible mechanisms of early molecular association and nucleation for amyloid generation. A wide-spectrum application of the current approach was also indicated by showing a successful utilization for ensemble structures of folded proteins. We propose that multiple linear regression in combination to experimental results will be highly promising for studies on protein misfolding diseases and functions by providing a convincing template structure.

**Graphic abstract:** 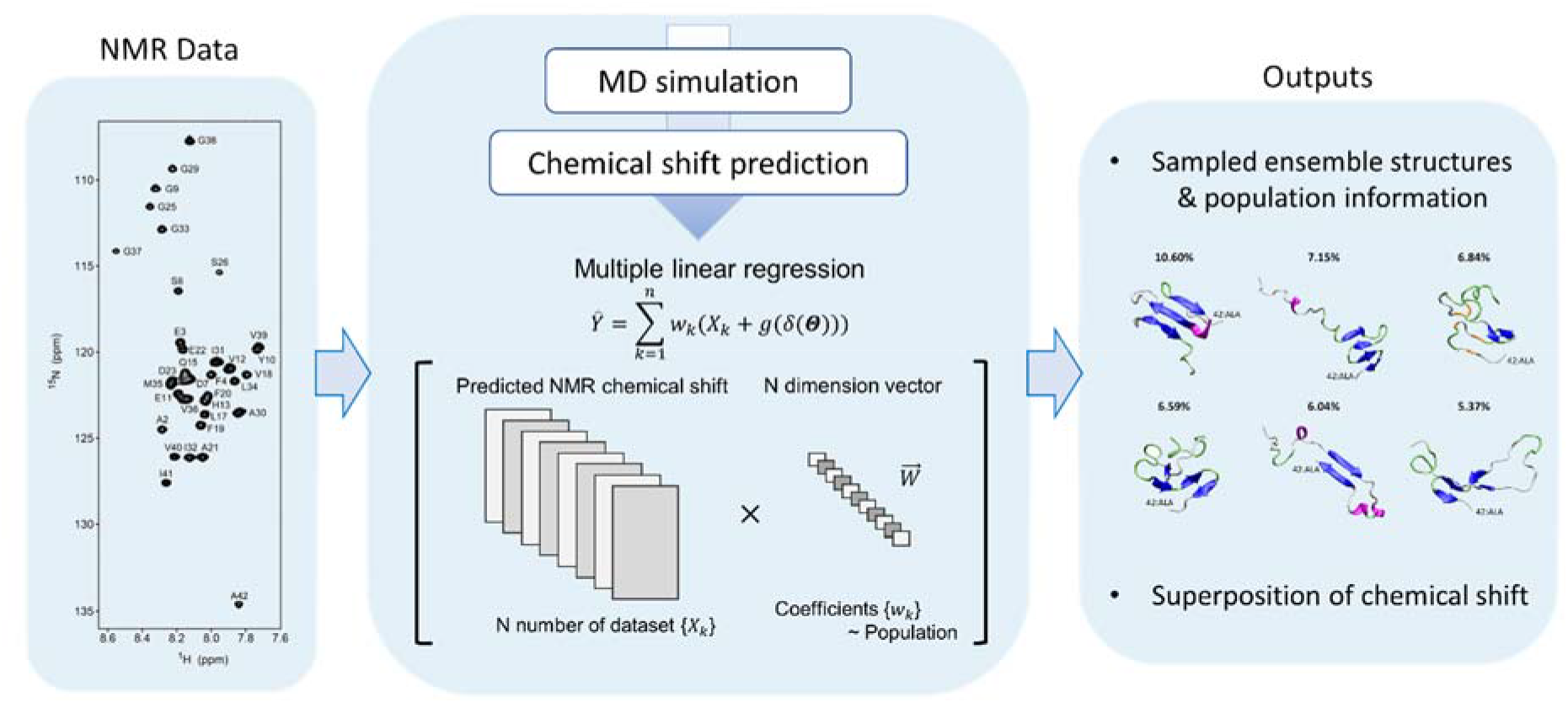

## Introduction

Proteins are a key player for numerous biological processes by using their unique three-dimensional or intrinsically disordered structures. Highly regulated intermolecular interactions endow proteins with the gain-of-function. However, interprotein mis-regulation often causes the loss-of-function and the gain-of-toxic function such as protein misfolding diseases due to aggregation^1–2^. It has been widely accepted that protein aggregation is deeply related to the onset and progress of various degenerative diseases^1–3^.

Several forms of aggregates of amyloid proteins such as amorphous aggregates, oligomers, protofibrils, and amyloid fibrils have been reported. Results revealed that morphology, structure, aggregation process, and cytotoxicity depend on the type of aggregates. Oligomers have attracted particular attention owing to their strong cellular cytotoxicity. Amyloid fibrils are best studied among protein aggregates. Unique properties of amyloid fibrils including cell-to-cell propagation, seeding capability, and prion-like behaviors are major risk factors for cellular homeostasis.

β-structure-rich amyloid fibrils largely display a two-step formation, slow nucleation producing a lag time and subsequent rapid elongation^4–6^. In contrast to nucleation-dependent amyloid formation, amorphous aggregation, protofibrillation, and oligomerizaion have been regarded as a nucleation-independent one-step process. Usually, they form instantly without an appreciable lag phase, manifest a growth phase, and, sometimes, aggregate between themselves. Of note, depending on experimental conditions, oligomers appear prior to amyloid formation as an intermediate, *i.e*. on-pathway oligomers, or remain as a dead-end product from the amyloid generation, *i.e*. off-pathway oligomers^7–8^. Disappearance of oligomers and subsequent productive nucleation for amyloid generation indicated that oligomers are kinetically stable and amyloid fibrils are thermodynamically stable aggregates as nationalized by Ostwald’s rule^9^.

A number of studies in the last two decades have provided fundamental information on protein aggregation using various experimental approaches including nuclear magnetic resonance (NMR), circular dichroism (CD), and fluorescence spectroscopies; however, much remains to be fully characterized to elucidate detailed mechanisms of the aggregation process. Especially, intermolecular interactions between soluble precursor proteins as well as their self-association and aggregation in the early stages of amyloid formation are fairly difficult to be examined at atomic or residue levels using direct experimental methods. Heterogeneous and transient initial structures and intermolecular interactions of disordered precursors in ensembles significantly hamper accurate and precise investigations on the early event of amyloid formation and the oligomerization. In order to overcome these difficulties, various *in silico* approaches including the molecular dynamics (MD) simulation and theoretical computation have been efficiently developed and exploited, which provided invaluable insights into the possible structural state and molecular association of amyloid proteins for aggregation^10–27^.

To date, numerous *in silico* investigations have been devoted to elucidating the structural properties of intrinsically disordered amyloid β (Aβ) peptides, of which aggregation in brains is suggested to be crucial for the pathogenesis of Alzheimer’s disease (AD) by inducing the neuronal cell death^10^. Previous studies reported that the formation of a β-sheet structure in the C-terminal region of Aβ (*i.e*., β-strands at positions 31–34 and 38–41) facilitates oligomerization and amyloid fibrillation^11–15^. N-terminal region, in contrast, is more likely to reduce the tendency of the formation of neurotoxic β-hairpin structures through direct interactions with the central hydrophobic region^16^. In addition to structural characterization, molecular actions of disordered Aβ monomers have also been explored in other complex molecular systems: 1) small molecule-Aβ interactions; 2) biomolecules/membranes-Aβ interactions; 3) spontaneous amyloid formation, etc^17–23^.

However, computational investigations on accurate structures of Aβ monomers which are physiologically disordered are still major challenges. A number of *in silico* studies used highly helical structures of Aβ monomers determined in water-alcohol mixtures with NMR spectroscopy as a starting structure even in aqueous solutions^24–25^. Moreover, disordered structures of Aβ monomers generated from highly helical structures were also often used as an initial template for simulations without experimental validation. As the starting structure is critical for both MD simulations and aggregation pathways^26^), much careful attention should be required for whether the initial structures of Aβ used in the simulation are sufficiently reliable.

Although *in silico* studies have remarkably improved our understanding of the monomeric structure and the molecular association (*e.g*., dimerization^12, 27^) and aggregation of Aβ, the data accuracy and reliability are always questionable due to the sensitivity of c omputational simulations to physical models. Thus, the incorporation of experimental d ata to computation is of particular importance to guide the rational design of molecular systems and validate *in silico* outcomes. In these regards, more reliable information on initial structures of Aβ monomers in an ensemble is required based on both *in silico* computation and *in vitro* experiments. Here, we examined initial structures of Aβ42 monomers in an ensemble in combination of computational and experimental approaches. Enormous ensemble structures of Aβ42 monomers were sufficiently sampled using the temperature replica-exchange MD (REMD) with an all-atomic protein model. Compelling initial structures in the ensemble best fit to experimental NMR chemical shifts with a novel multiple linear regression were successfully revealed (**Fig. 1**). Aβ42 ensemble structures currently elucidated well explained possible mechanisms underlying the molecular association and aggregation for early-stage Aβ42 amyloidogenesis. Our new hybrid approach will contribute to obtaining the insights into the molecular mechanisms of the Aβ42-related aggregation and the development of therapeutic molecules targeting AD. It will be also useful for the structural and functional studies on intrinsically-disordered proteins.

**Figure 1.**
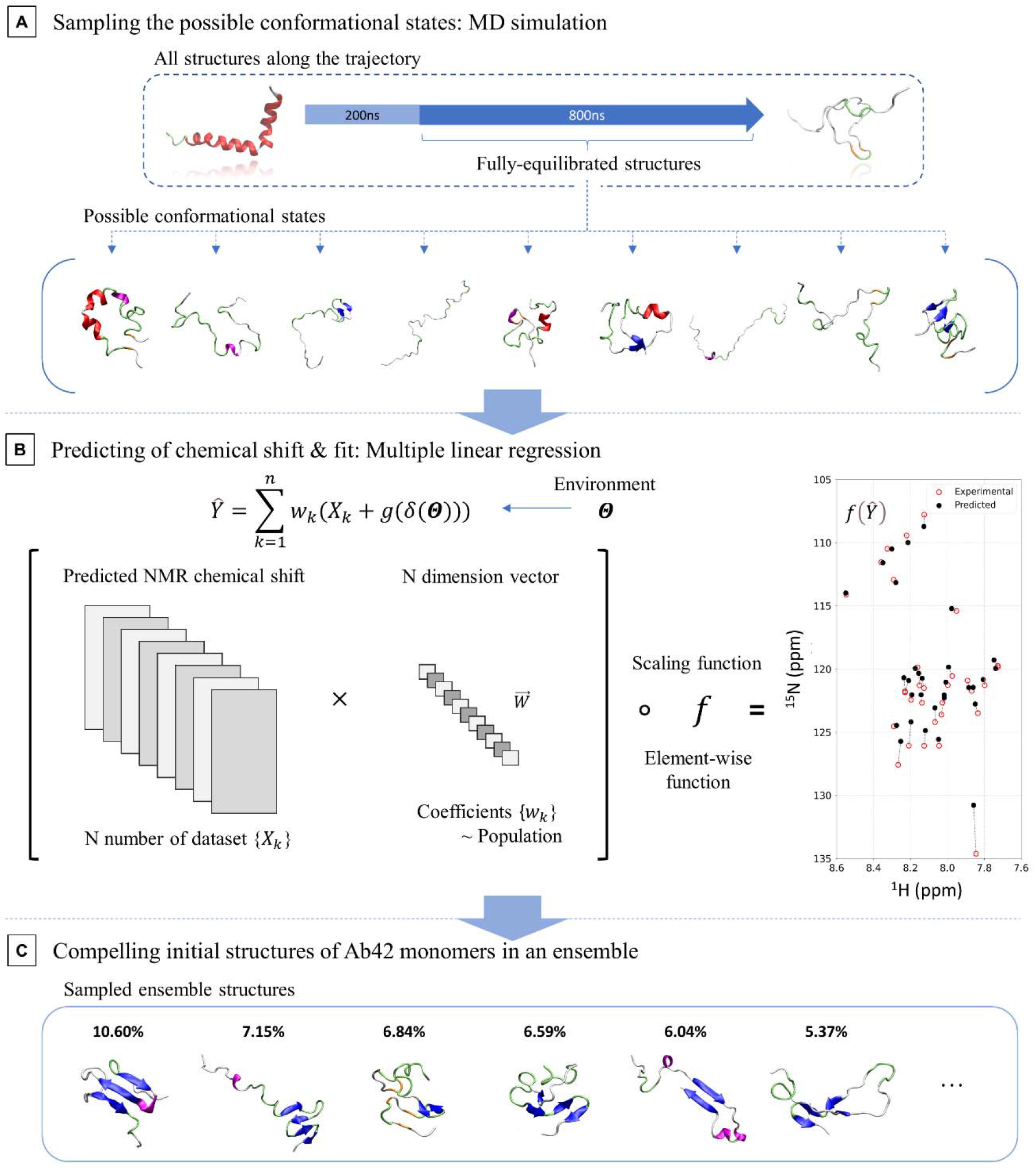
Schematic flow of the multiple linear regression approach. Three main steps are sequentially shown from A to C. (A) Possible Aβ42 monomer structures obtained from replica exchange MD simulation (REMD). (B) Best-fit of predicted NMR chemical shifts obtained using REMD and UCBSHIFT to the most compelling ensemble of initial Aβ42 structures using the multiple linear regression. (C) Solution structures of Aβ42 monomers in the ensemble (*i.e*., sampled ensemble structures).

## Results

### Characterization of averaged Aβ42 conformations in solution using biophysical approaches

In order to reveal an averaged conformational state of Aβ42 monomers at a physiological pH condition, we performed colloidal and structural characterization of Aβ42 using several biophysical approaches including dynamic light scattering (DLS) measurements as well as CD and solution NMR spectroscopies at pH 7.5 (**Fig. 2**) We first tested a colloidal state of Aβ42 molecules using the size distribution obtained by DLS. As shown in **Fig. 2A**, monodispersed DLS peaks centered at the hydrodynamic radius (*R_H_*) of ~1.666 nm, similar to *R_H_* of ~1.6 nm obtained by Förster resonance energy transfer (FRET) and fluorescence correlation spectroscopy^28^, were detected without any additional peaks of large *R_H_* values. This indicated that Aβ42 was in a monomeric state without aggregates such as oligomers and amyloid fibrils.

**Figure 2.**
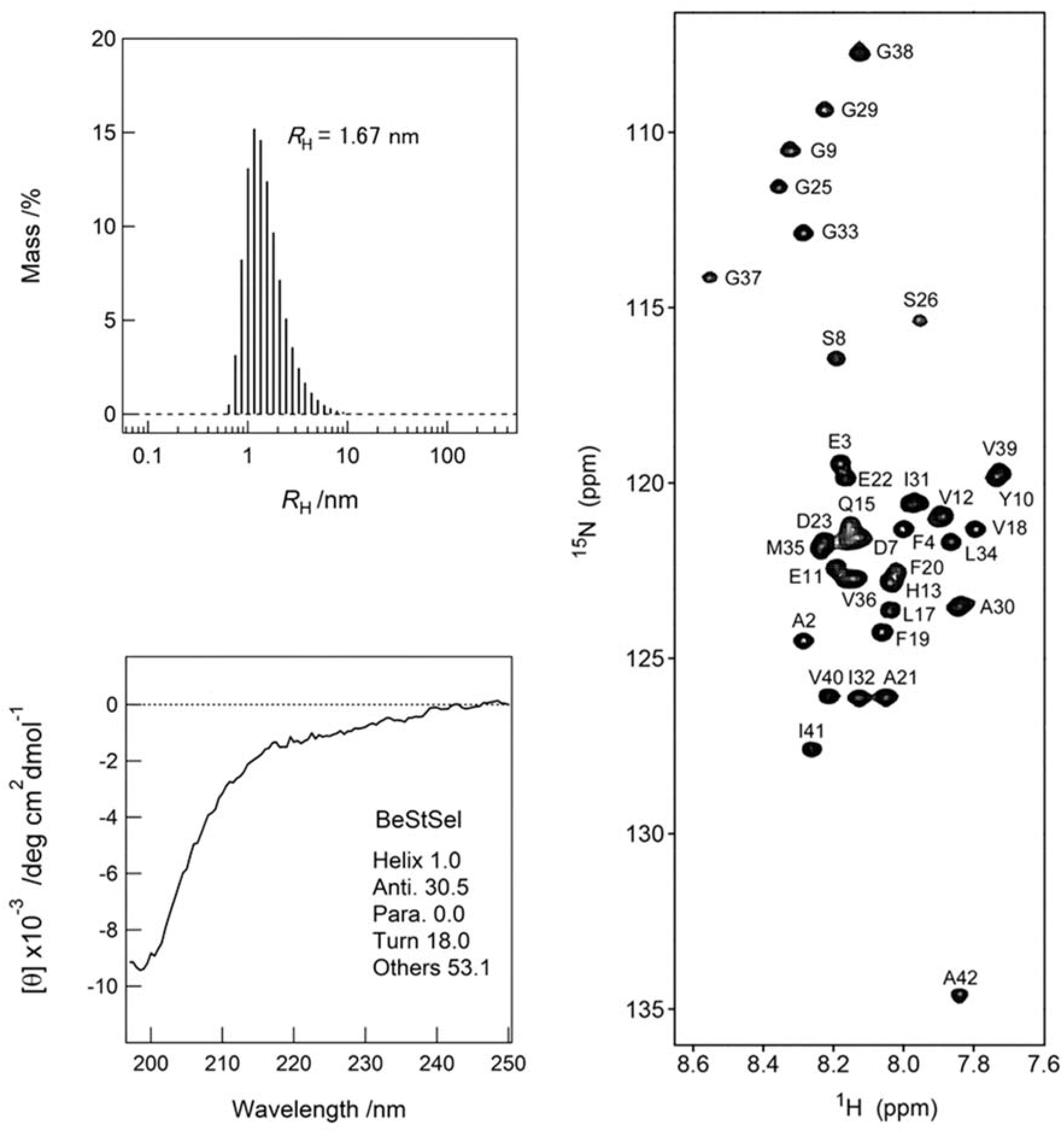
Structural and colloidal characterization of Aβ42. (A-C) Several biophysical approaches were used: dynamics light scattering (A), far-UV circular dichroism spectroscopy (B), and two-dimensional solution NMR (SOFAST-HMQC) spectrum (C). *R*_H_ in A indicates the hydrodynamic radius. The contents of the secondary structure calculated using the BeStSel are shown in B.

Next, the secondary structures of Aβ42 were examined using the far-UV CD spectroscopy and an algorithm which calculates the content of the secondary structure.^29^ The far-UV CD spectrum of Aβ42 showed a single negative peak at ~200 nm with a weak characteristic band in the region between 210 to 230 nm (**Fig. 2B**), suggesting that secondary structures of Aβ42 monomers were highly disordered as reported in previous studies.^30–32^ For further characterization, we determined the proportion of the secondary structure using BeStSel^33^: the contents of the secondary structure were calculated to be ~1.0% of α-helix, ~30.5% of anti-parallel β-sheet, ~18.0% of turn, and ~53.1% of others mainly including random coil-like structures.

The two-dimensional (2D) ^1^H-^15^N band-selective optimized flip-angle short transient (SOFAST)-heteronuclear multiple quantum coherence (HMQC) spectrum provided atomic level evidences of largely unfolded conformations of Aβ42 monomer (**Fig. 2C**). The SOFAST-HMQC spectrum of Aβ42 obviously presented a narrow distribution of chemical shifts of amide protons spanning from ~7.6 to ~8.5 nm as well as intense NMR peak intensities, which are well-known characteristics of unfolded proteins.^34^ Taken together, our data demonstrate that Aβ42 initially exists as a monomer and its averaged structure is largely disordered with partial β-structures.

### Comparison between replica-exchange MD simulation and experimental results

Previous MD simulation studies have been performed using both the conventional and enhanced sampling methods such as temperature replica exchange MD (REMD) in order to sample the locally stable structures which are generally difficult to be detected.^14, 35–36^ Thus, we first used the REMD sampling method in an attempt to fully explore the ensemble states of Aβ42 monomer in both the implicit and explicit solvent models after 200-ns trajectories (80,000 ensemble structures and the 1-μs simulation in total) (see supporting information).

Analyzed results revealed the distribution of the radius of gyration (*R*_g_) and the proportion of the secondary structure of Aβ42 monomers (**Fig. S1A, B**). The distribution of *R*_g_ obtained by the implicit solvent model was more widespread than that of the explicit solvent model (**Fig. S1A**). Average values of *R*_g_ for the explicit and implicit solvent models were 1.091 and 1.729 nm respectively, indicating that averaged Aβ42 conformations in the explicit solvent model are more compact than those in the implicit solvent model. By using empirical relationships between *R*_g_, *R*_H_, and the residue number of intrinsically disordered proteins (IDPs),^37^ *R*_H_ values in the explicit and implicit solvent models were calculated to be 1.533 and 1.835 nm, respectively, which did not match the *R*_H_ value of ~1.666 nm directly obtained by the DLS experiment. In addition, the proportion of β-structures computed based on the two solvent models was also inconsistent with the calculated proportion using the BeStSel and experimental results of the far-UV CD spectra (**Fig. S1B**). The averaged secondary proportion shows 7.22%, 3.31% α-helix and 6.33%, 4.56% β-sheet in explicit and implicit solvent model, respectively. However, the BeStSel result shows more less α-helix proportion (1.0%) and larger β-sheet proportion (30.5%) than averaged results.

We further generated a series of NMR chemical shift prediction data from vast MD simulation trajectories using the several algorithms of SHIFTX2^38^, SPARTA+^39^, and UCBSHIFT^40^ (**Fig. S1**). We obtained a total of 80,000 PDB inputs from 200 to 1000 ns for each implicit and explicit solvent MD simulation. SHIFTX2 and SPARTA+ have been generally used for globular proteins, by comparison, UCBSHIFT is a newly developed algorithm to predict the NMR chemical shift of both globular and disordered proteins like IDPs. We compared the average of chemical shifts computed using UCBSHIFT (**Fig. S1C, D and Table S1**), SHIFTX2, and SPARTA+ (**Table S1**) with experimental NMR chemical shifts (**Fig. 2C**). All Score_total_s, denoted *R^2^* scores which represent the proportion of variances for the chemical shifts, of averaged chemical shifts are negative or small values (details of scoring method in supporting information).

Although narrow distributions of chemical shifts in NMR spectra obtained by both the computation and direct NMR experiments appeared to be similar, almost all of the chemical shifts calculated using MD results did not match those obtained by NMR experiments. Collectively, all these comparative results indicated that ensemble conformations of whole Aβ42 monomers obtained from *in silico* MD simulations do not convincingly explain *in vitro* experimental results.

### Best-fit ensemble structures of Aβ42 monomer from the linear regression approach

As the above-mentioned computation did not provide reasonable results, we set out to develop a regression method which bridge *in silico* and *in vitro* experimental results. Our hypothesis is that MD simulations with an enhanced sampling method can explore sufficiently possible ensemble states of Aβ42 monomer, and, thus, a NMR chemical shift pool for whole Aβ42 is predictable. If a multiple linear regression approach is valid, it will find the most compelling ensemble (hereafter referred to as “sampled ensemble”) which consists of initial Aβ42 structures (hereafter “sampled ensemble structure”) by fitting NMR chemical shifts predicted to those experimentally obtained. Thus, individual Aβ42 monomeric structures in the ensemble are determined with and their populations (**Fig. 1**). Our multiple linear regression method is specialized for finding the most reasonable coefficient (*w_k_*) in given ensemble, which represents the probability of each conformational state. In this case, the reasonable coefficients should satisfy the following three conditions: 1) the sum of coefficients is equal to 1, 2) each coefficient has a positive value, and 3) multiple linear regression does not possess interception.

We performed the multiple linear regression for all sets of predicted NMR chemical shifts obtained using the MD trajectory and three algorithms, SPARTA+ (**Fig. S2A, B**), SHIFTX2 (**Fig. S2C, D**), and UCBSHIFT in both solvent models (**Fig. 1**). For the more reliable multiple linear regression analyses, a scaling function to minimize the sum of squares of residual errors for the chemical shift of ^1^H and ^15^N was introduced. We also developed an algorithm to revise the effectiveness in the ^1^H chemical shift which is often caused by experimental conditions such as temperature using the non-negative least squares regression and repeated optimization to be *w_k_* of 1 (details in supporting information). All regression scores of the multiple linear regression are represented in supporting information (**Tables S2**). All Score_total_s after the regression are closed to 1, which suggests that the regression provides more well-fitted ensemble than averaged chemical shift.

Among three analyzed results, the multiple linear regression with UCBSHIFT manifested the highest regression scores with the most similar NMR chemical shifts to experimental one, indicating that the UCBSHIFT-based multiple linear regression analysis reflect experimental results more faithfully in both solvent models (**Fig. 3B, D**). It is thought that UCBSHIFT offers best prediction as it was tailored algorithms for IDP-like proteins. Top 6 major structures (**Fig. 3A, C**) and top 10 major structures (**Fig. S3**) of Aβ42 monomer in the regression analyses with UCBSHIFT were presented. In addition, we obtained a similar structural ensemble of Aβ42 based on published chemical shift data (BMRB Entry 25218^33^) (**Figs. S4–S5**), which is left-shifted approximately 0.2 ppm in the ^1^H axis compared to our chemical shift due to distinct experimental temperature. This spectra shift depends on increasing temperature for Aβ40 is already reported in previous study^41^. Similar illustration about major structures of Aβ42 monomers from other chemical shift prediction algorithm is also in supplementary data. (SPARTA+; **Figs S6–S7** and SHIFTX2; **Figs S8–S9**)

**Figure 3.**
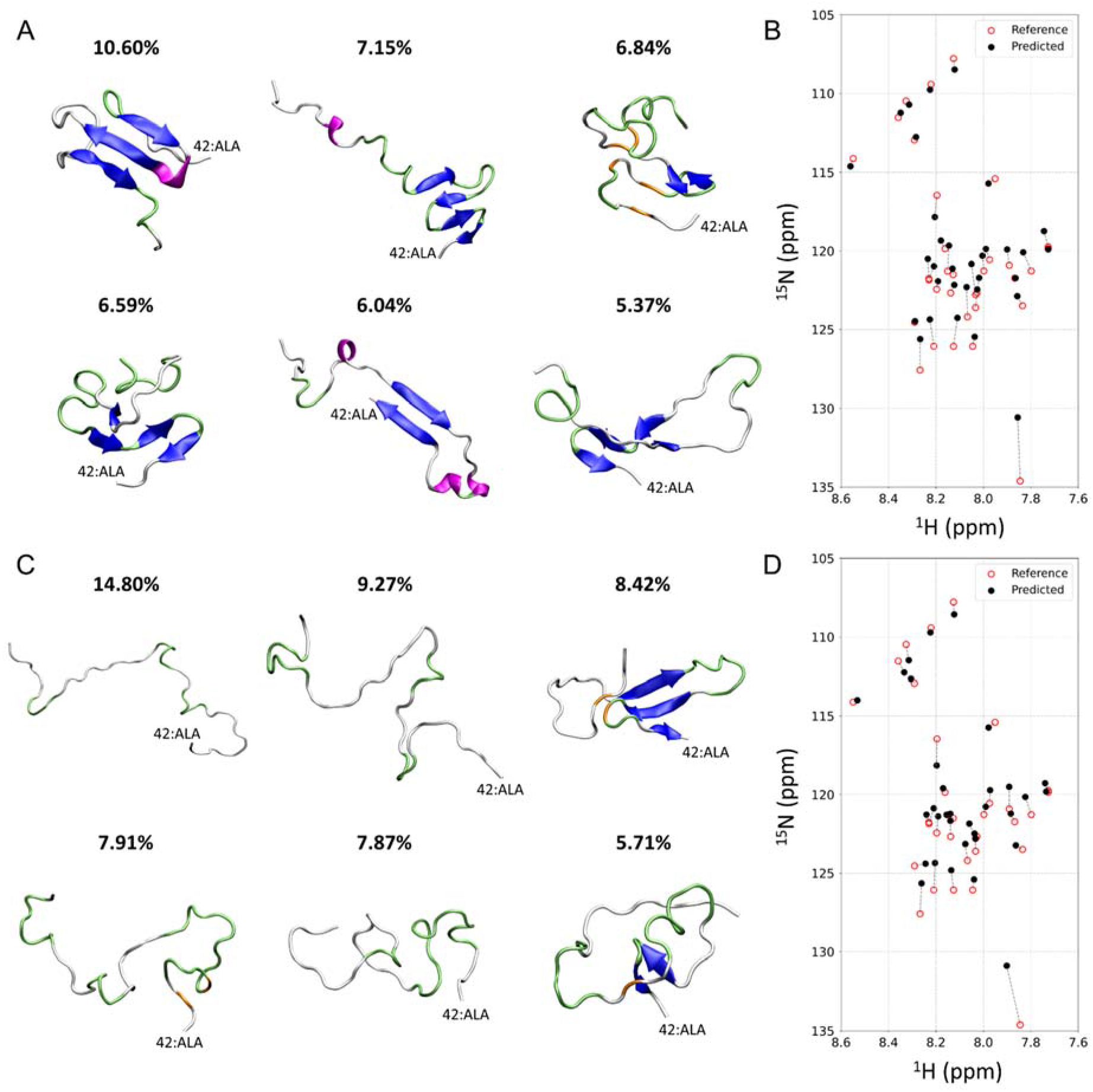
The major structures of Aβ42 monomers in the ensemble. (A, C) Representative Aβ42 structures with higher populations obtained using the experimental NMR chemical shift and UCBSHIFT in the explicit (A) and implicit solvent model (C). Secondary structures are colored as follows: α-helix (red), β-sheet (blue), 3_10_-helix (magenta), β-bridge (orange), turn (lime), and coil (white). The weighted average of chemical shift data by the proportion in the explicit (B) and implicit solvent model (D) is superimposed with experimentally-determined chemical shifts.

### Validation of the regression approach with comparison to experimental results

In addition to verification of the NMR chemical shift above, we further evaluated our regression approach based on the experimental results, the hydrodynamic property and the proportion of the secondary structure of the conformations of Aβ42 ensemble in solution (**Fig. 4**). The weighted averages of *R*_g_ for sampled ensemble structures were first computed based on the regression coefficients. Then, we quantified a variance of the prediction results using each che mical shift prediction as a discrete random variable of the conformational space (**Figs. 4A and S10A**). The weighted averages of *R*_g_ in the explicit and implicit solvent model with UCBSHIFT are 1.179 nm and 1.631 nm, respectively. *R*_H_ was further calculated to be 1.586 nm and 1.799 nm, respectively, using the same relationship used earlier.^37^ These values are more simi lar to the experimental *R*_H_ value of ~1.666 nm than that obtained by REMD without the linear regression (**Fig. 2A**). Thus, it was indicated that our regression approach is more appropriate to demonstrate Aβ42 structures in an ensemble.

**Figure 4.**
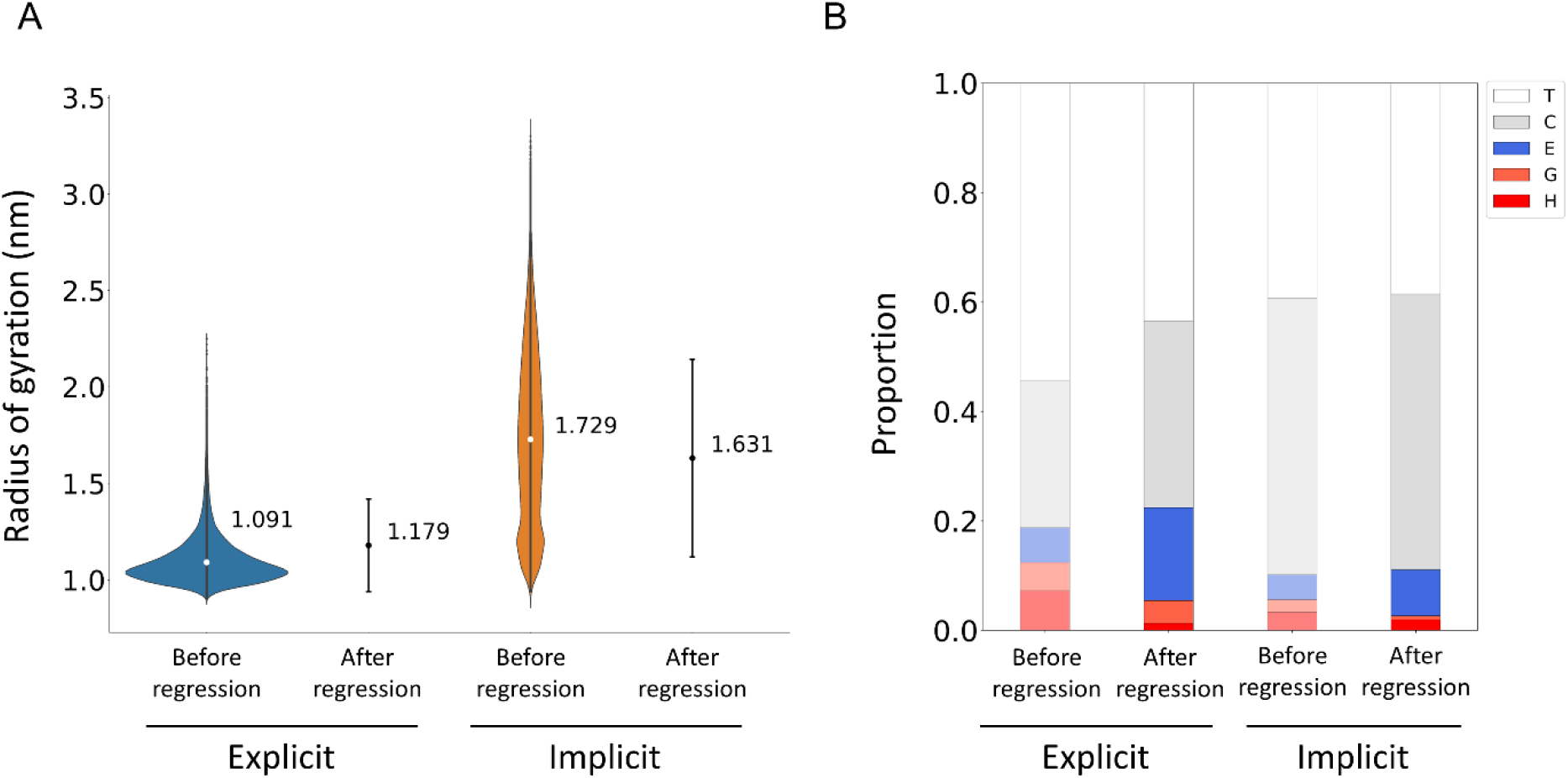
Effects of linear regression on structural properties of Aβ42 monomers in ensembles. (A, B) The radius of gyration (*R_g_*) (A) and proportion of the secondary structure (B) of Aβ42 before (Explicit and Implicit) and after multiple linear regression (UCBSHIFT_exp_ and UCBSHIFT_imp_). Error bars in A indicate the standard deviation. The type of the secondary structure is shown with the single letter and color as follows: H, α-helix (red); G, 3_10_-helix (orange); E, β-sheet (blue); C, coil (gray); and T, turn (white).

Next, we calculated and analyzed the secondary structures of Aβ42 structures in the sample ensemble. Secondary structures of the sampled ensemble showed increased portions of β-sheet formation in both implicit and explicit solvent models with UCBSHIFT, SPARTA+, and SHIFTX2 (**Figs. 4B and S10B**). Before regression with UCBSHIFT, the averaged proportions of the secondary structure of α-helix, 3_10_-helix, β-sheet, coil, and turn were 7.22%, 5.20%, 6.33%, 26.80%, and 54.45% in the explicit solvent model, and 3.31%, 2.29%, 4.56%, 50.54%, and 39.30% in the implicit solvent model, respectively. After regression with UCBSHIFT, the proportions were 1.30%, 4.14% 16.97%, 34.17%, and 43.42% in the explicit solvent model, and 1.96%, 0.68%, 8.43%, 50.34%, and 38.59% in the implicit solvent model, respectively. Importantly, sampled ensemble structures of Aβ42 monomers, *i.e*., results with both solvent models showed more similar secondary structures to those predicted using CD spectra (**Fig. 2B**). A number of compact structures in the explicit solvent model were detected. Hence, we conceive that a higher tendency of the explicit solvent model to sample β-sheet formation may be attributable to an easier conformational collapse compared to the implicit solvent model.

Considering all the data for regression analysis comprehensively, the regression results using UCBSHIFT provides the best score among all prediction algorithms. It is probably because UCBSHIFT gives better prediction than other algorithms for IDP-like protein. And the sampled ensemble using explicit solvent model with UCBSHIFT prediction gives us better explanation for the experimental data than implicit solvent model. Therefore, afterwards, we focused on the regression results using UCBSHIFT with explicit solvent model to progress getting more structural information for Aβ42 protein. All of the data discussed later used data based on UCBSHIFT with explicit solvent model.

### Characterization of averaged structures of Aβ42 monomer in ensembles

Ensemble - of Aβ42 monomer in the explicit solvent model determined using UCBSHIFT were further analyzed at the residue level, and improvement of structural refinement by the multiple linear regression was assessed (**Fig. 5**). All Aβ monomers in ensembles before the multiple linear regression showed intrinsically disordered features without three-dimensional conformations (**Fig. 5A**). Turn and random coil structures were mainly observed with relatively minor β- and helical conformations. β-sheet formation was prominent in the central hydrophobic core (CHC) and C-terminal region than the other parts. Helical structures were exclusively formed in the C-terminal part with a higher proportion than β-structures.

**Figure 5.**
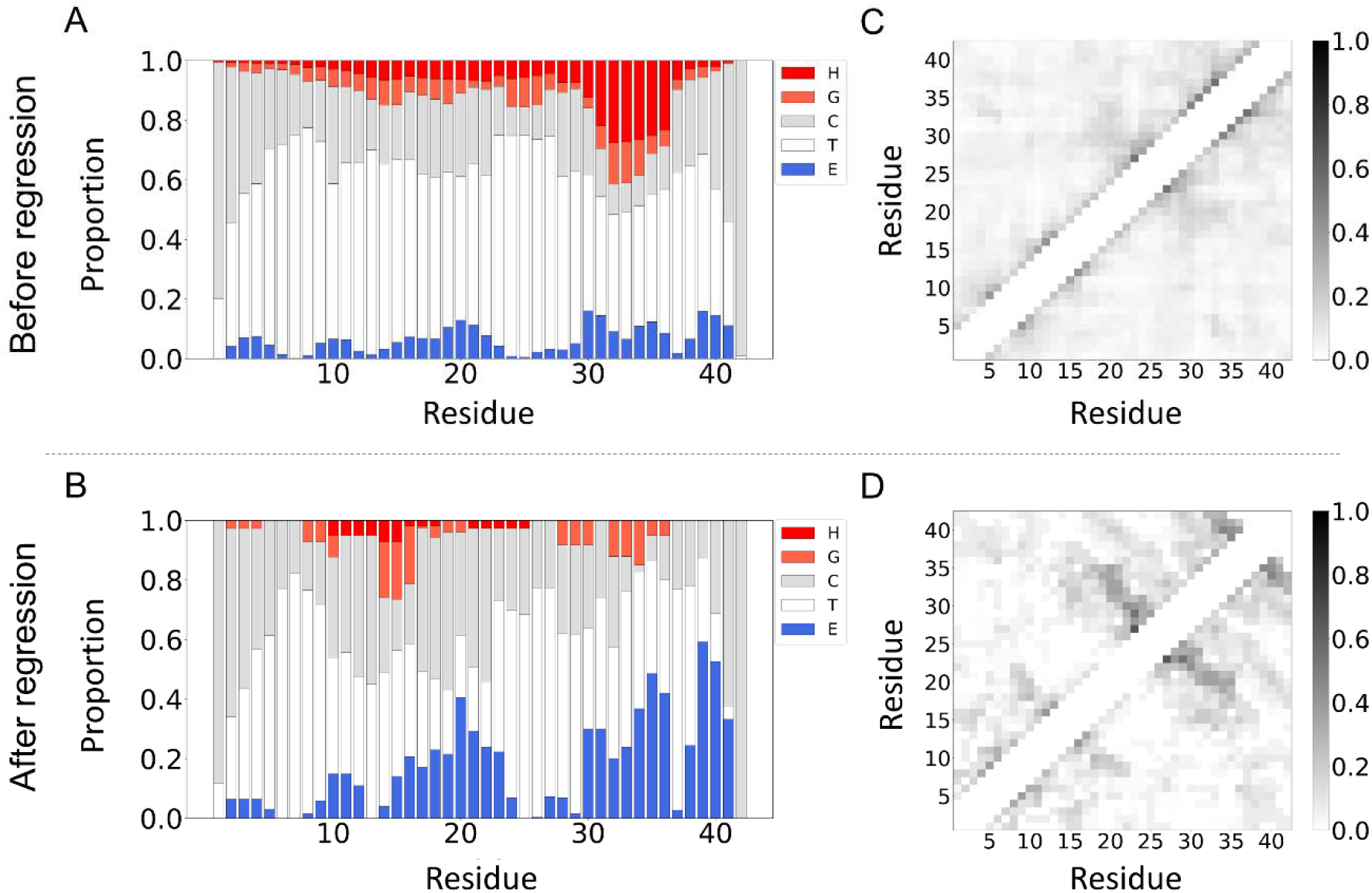
Linear regression-dependent changes in the secondary structure and intramolecular contact of Aβ42. Proportion of the secondary structure (A and B) and contact map (C and D) of Aβ42 monomer ensemble in the ensemble before (A and C) and after the regression approach (B and D) in the explicit solvent model. The type of the secondary structure in A and B is shown with the single letter and color as follows: H, α-helix (red); G, 3_10_-helix (orange); E, β-sheet (blue); C, coil (gray); and T, turn (white). Degree of the contact between residues of Aβ42 in C and D is scaled from 0.0 to 1.0 which is shown by the gradation (right).

Noticeable changes in the secondary structures of Aβ42 monomer were observed after the multiple linear regression by keeping intrinsically disordered features (**Fig. 5B)**. Sampled ensemble structures of Aβ42 increased the β-sheet propensity with decreases in helical contents. Proportions of β-structures increased throughout an Aβ42 monomer, and, several distinguishable patterns of local β-sheet formation were detected in the N-terminal (residues 2–6 and 10–14), CHC (residues 17–23), and C-terminal regions (residues 27–41). These results were consistent with CD-based analyses which showed appreciable contents of β-structures (~30%) (**Fig. 2B**). At the same time, tendencies of the formation of helical and turn structures were attenuated, and, a small proportion of helical structures appeared in the range of residues 10–16.

In order to examine further conformational features and impacts of the multiple linear regression, the contact maps of Aβ42 structures in ensembles before and after regression treatment were constructed (**Fig. 5C, D**). No significant patterns were obtained in the explicit solvent model before linear regression (**Fig. 5C**). However, the multiple linear regression obviously revealed several anti-parallel patterns in the sampled ensemble structures (**Fig. 5D**). Especially, an anti-parallel pattern between the residues 19–25 and the residues 27–34 and another anti-parallel pattern between the residues 32–36 and the residues 39–42 are strongly appeared in the contact map. These anti-parallel patterns in the contact map indicated the formation of several β-strands (**Fig. S3A**).

To verify these significant β-sheet patterns, we further examined the contact map of the implicit solvent model. Before the regression, there was a marked tendency of β-sheet formation in the CHC (residues 17-23) and C-terminal regions (residues 29-42) (**Fig. S11A, C**). Anti-parallel β-sheet patterns in the contact map obtained after the regression of the explicit solvent model were also broadly detectable in the contact map of the implicit solvent model (**Fig. S11B, D**). These findings suggested that the trend of the contact maps in both solvent models was similar although detailed contacts were not identical. Thus, the Aβ42 monomers in the sample ensemble have a high potentiality to contain more than two β-strands in our experimental data.

### Expandability of the linear regression approach to ensemble structures of folded proteins

To generalize and validate our mathematical methodology, we applied the multiple linear regression to two different folded proteins, ubiquitin and chymotrypsin inhibitor II (CI2). We used the implicit solvent model since it can fully explore major conformational states of folded proteins, and UCBSHIFT for the prediction of the chemical shift prediction.

Ubiquitin (PDB code: 1D3Z^42^) (BMRB Entry 15410^43^) is a small protein with 76 residues, and, exists in all eukaryotes. A unique but generalized “ubiquitin fold” is characterized by a complex topology consisting of a 5-stranded β-sheet, an α-helix, and a short 3_10_-helix.^44^ Together with these properties and several advantages such as no prosthetic groups and disulfide bonds as well as high solubility and thermostability,^44^ it has been extensively used as a model protein for the studies on folding, structure, and NMR. We ran a 1-μs REMD simulation to maximally explore possible conformational space, and performed the regression approach with the optimization process as in the case of Aβ42. (Figs. 6A and S12). Detailed scores of chemical shift values are summarized is in Table S3–S4. Of note, results unveiled that all structures of the sampled ensemble were similar to solution NMR structures in PDB (Fig. 6B).^42^ CI2 (PDB code: 3CI2^45^) (BMRB Entry 4974^46^) was further examined using the same procedures. CI2 is a small 64-residue serine proteinase, and has 4 β-sheet structures and an α-helix.^46–47^ Although the chemical shift prediction result was not perfectly fit to reference data, all structures in the sampled ensemble were similar to NMR ensemble structures previously reported (Figs. S13–S14 and Tables S5–S6)^45^, suggesting that our approach is still valid for CI2. The comparisons between ensemble structures obtained using our linear regression approach and NMR spectroscopy are shown for both ubiquitin and CI2 (Fig. S13B). Overall, all these results verify that our multiple linear regression method is not limited to unstructured Aβ42, and, also valid for the investigation of structural ensembles of folded proteins.

**Figure 6.**
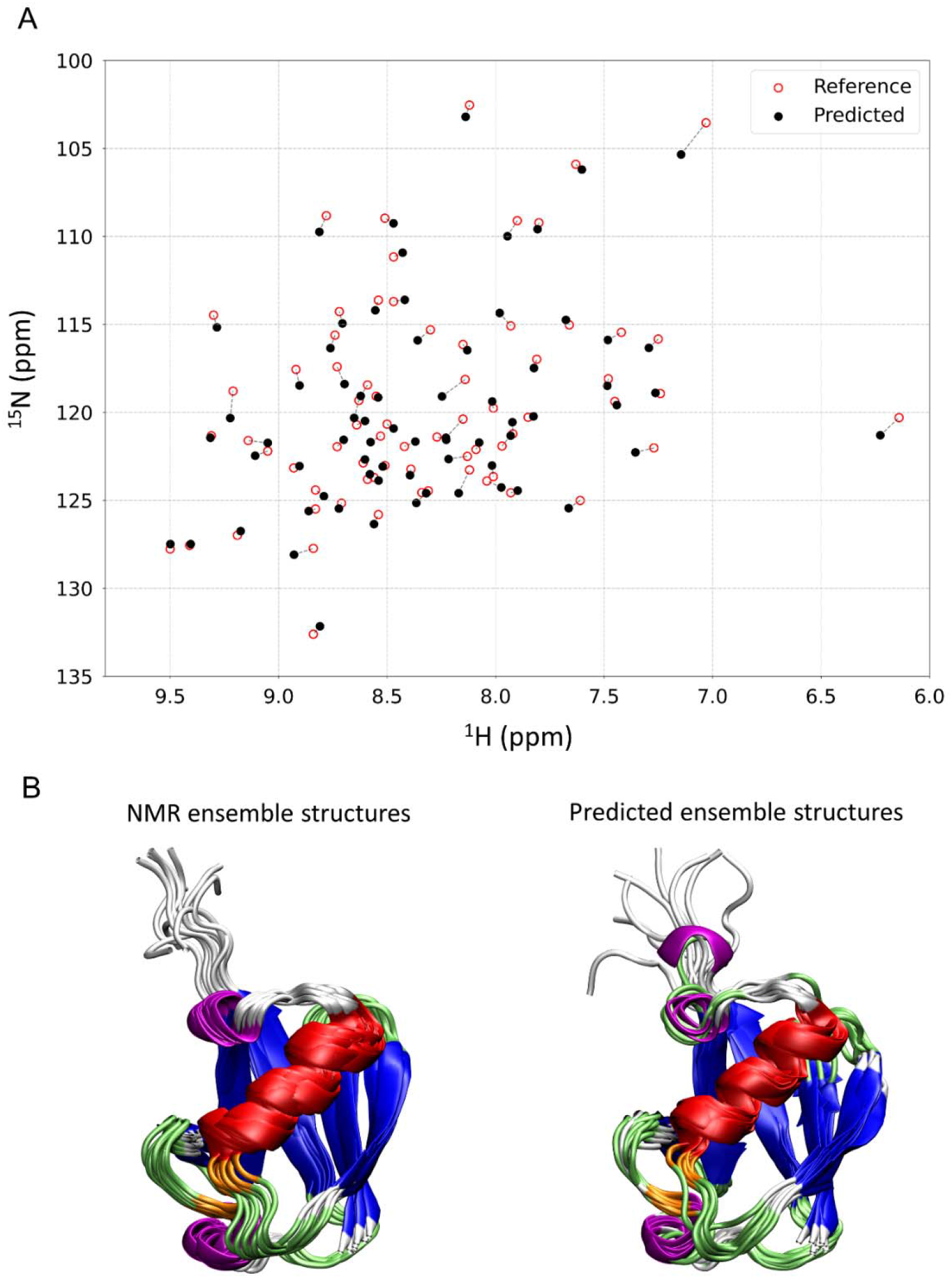
Multiple linear regression approach for ubiquitin. (A) Comparison of chemical shifts calculated using linear regression (red) with those directly obtained using NMR spectroscopy (black) (BMRB Entry 15410). (B and C) The contact map (B) and ensemble structures (C) of ubiquitin after multiple regression are shown. The secondary structures are colored as follows: α-helix (red), β-sheet (blue), 3_10_-helix (magenta), β-bridge (orange), turn (lime), and coil (white).

## Discussion

The overall mechanism underlying Aβ aggregation has been increasingly revealed; however, early-stage intermolecular interactions and associations of Aβ monomers for the formation of oligomers and amyloid fibril still remain elusive. In this study, we hybrided *in vivo* and *in silico* results to compensate demerits and utilize advantages of two different methods, and, finally generated a new methodological approach which reveals heterogeneous initial structures of Aβ42 monomers in the ensemble. Accordingly, we exclusively describe several representative initial structures of Aβ42 monomers which would be important for understanding key of Aβ aggregation.

Aβ protein is easy to aggregate in room temperature. This aggregation and accumulation of Aβ protein cause AD. Because of this pathogenetic necessity, it is important to study Aβ aggregation. Although A*β* protein have no well-defined secondary structure, the ensemble of protein in specific environment frequently reveals several local secondary structures. These patterns can be appeared in proportion to its energetic states. In this situation, some specific structures of ensemble provide additional opportunities to aggregate as nucleus. Knowing structures in an ensemble is important to have insights into the molecular mechanism of the polymorphs of amyloid fibrillation at the early event. Depending on ambient conditions, major and minor populations will be changed in new equilibrium, and, in turn, resulting in the formation of a new nucleus for amyloid formation. This view is based on thermodynamics. But, it would kinetically possible that a structure with a minor population can form rapidly a nucleus, and, immediately generate short amyloid fibrils. Although the population of aggregation-prone structures is minor, aggregation event is sequentially spread out from the small number of initial structures.

Recent studies paid attention to a variety of self-association modes of Aβ.^48–49^ Tycko^48^ reported that Aβ can construct varied specific fibril morphologies. The polymorphic characteristics can define a molecular mechanism for about different range of toxicity. It suggests that specific environment affects to make “the initial seed” of a specific aggregation for Aβ. A lot of previous studies investigated the aggregation condition of Aβ using NMR and further analysis.^48–51^ Similar initial structure-dependent aggregations were also reported in other proteins such as cytochrome *c*^52^, tau^6^, α-synuclein^53^ and β2-microglobulin.^54^ For example, membrane binding of α-synuclein induces the membrane-unbound regions to be relatively unstructured so that this membrane disruption effect makes different aggregation patterns of α-synuclein.

We suggested representative local β-sheet structures of Aβ42 ensemble. These local formations are supported by the CD spectra result in the region between 210 to 230 nm. These local β-sheet structures are influenced by the central hydrophobic core (CHC) and hydrophobic C-terminus region of Aβ42.^55–56^ Several MD simulation studies suggested that Aβ42 forms a major anti-parallel β-hairpin involving the central hydrophobic cluster residues 16–21 with residues 29–36.^15, 57^ Turn 26–27 occurs simultaneously with β-sheet structure involving the central hydrophobic cluster residues 16–21 with residues 29–36.^57^ Our results showed a similar appearance which has less secondary structures at the range of residues 25–28 (**Figs. 5B and 7**) The structure which has the most highest occurrence (1^st^ structure of figure 3A) in regression result are proximal to these tendency (**Fig. 3A**). These Aβ monomer structure might be energetically stable so that it appears more frequently. From a lot of studies, the CHC region and C-terminus region of Aβ42 seem to have a crucial role to compactness of Aβ42 for all conformations including monomer, dimer, oligomer and furthermore fibril state.^58–60^ Intra/inter-molecular hydrophobic interactions between these regions control the amyloid-formation pathways of Aβ42.^58, 60^ Considering these evidences comprehensively, the most highest occurrence structure has a possibility to be initial structure of aggregation such as nucleation-dependent amyloid formation including oligomerization and surface-catalyzed secondary nucleation and nucleation-independent elongation. The general kinetic model of amyloid fibril formation was reported on previous studies.^11, 61^

**Figure 7.**
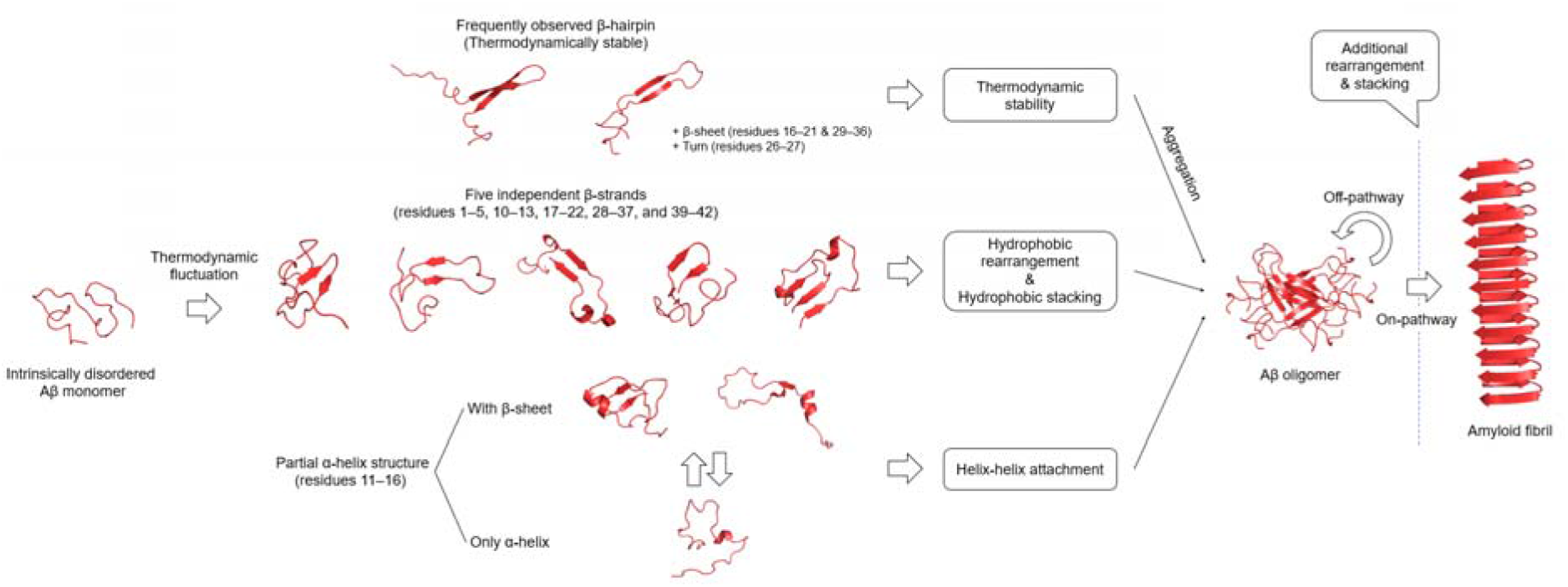
Possible aggregation mechanism from generated Aβ42 ensemble.

Nucleation-independent oligomerization of Aβ protein is commonly affected by certain hydrophobic units. Some previous studies reported the importance of extended β-hairpins to form β-sheet oligomers.^54, 62–63^ And, Yang and Teplow^13^ reported the five independent β-strands, comprising residues 1–5, 10–13, 17–22, 28–37, and 39–42. These folding units are also shown in Aβ42 dimer simulation.^62^ Also, Hoyer et al^64^ suggested the β-hairpin structure with several hydrophobic clusters (L17, F19, I32, L34, and V36) constitutes an intermediate conformation on the pathway to amyloid fibrils for Aβ40 protein. The Aβ40 oligomers might form by hydrophobic stacking of β-hairpins.^64^ Our residual secondary proportion result for sampled structures shows similar significant local β-sheet patterns (**Figs. 5B and 7**). The local β-sheet distribution can be distinguishable into small number of residues 2–5, 9–12, wide range of CHC region which consists of residues 15–23, and two separated hydrophobic C-terminus region spanning residues 30–36 and 38–41. Although these patterns are not appeared simultaneously, these characteristics of Aβ42 may make to progress into dimerization and oligomerization. Furthermore, the oligomerized Aβ protein can be continued to protofibril formations by hydrophobic stacking.

Elongation of amyloid fibrils and protofibrils in growth phase is nucleation-dependent amyloid formation because their fibral shapes depend on the initial nucleus of early-phase. Karamanos et al^54^ reported the rearrangements of the hydrophobic core plays key role in aggregation, which results in exposure of buried hydrophobic residues. A native-like arrangement of the main-chain is enough to expose hydrophobic parts of the protein that presumably act as nucleation points for amyloid aggregation. Also, they mentioned unfolding energies, hydrophobicity and propensity of inter/intra-molecular contacts are linked to amyloid propensity. Previous MD strudies^13, 65^ reported that the C-terminus of Aβ42 protein has an frequent β-sheet forming tendency. This thermodynamic feature makes inter-molecular contacts in elongation phase and sequential rearrangements of the hydrophobic core might be extended to amyloid elongation. Another previous solution NMR study showed the residue-specific solvent protection map within the Aβ42 fibril.^66^ It is composed to two protected regions, spanning residues E11-G25 and K28-A42. These hydrophobically protected regions can be assembled from rearrangements of the hydrophobic cores, comprising residues 10–13, 17–22, 28–37, and 39–42. The partially β-sheet forming structures are shown in supplementary figure S3A.

Another question is that it is possible to understand the primary nucleation event from our results. In the primary nucleation, a partial α-helical structure is important intermediated state for nucleation-dependent amyloid formation.^26^ Lin et al^26^ suggested various aggregation pathway of Aβ40. According to their aggregation pathway with conceptual energy diagrams, partial α-helical structures are aggregation-prone and sticky because of their enhanced hydrophobicity. These partial α-helical structures might strengthen the helix–helix interaction, causing rapid aggregation in primary nucleation. The partial α-helical structure is revealed on the residues E11, V12, H13, H14, Q15, and K16. Our residual secondary proportion data supports this significant composition in E11 to K16 region, which does not have only α-helix, but also 3-10 helix, too. Our sampled representative structures show a reliable partial α-helical structure (7th structure of Figure S3A) which satisfies previous thermodynamic feature of C-terminus.^65^ However, we did not ensure that this structure to be able to initial seed for primary nucleation and the measured experimental chemical shift which is used in performing multiple linear regression shows oligomerization process in progress. If we can measure sequential time series NMR chemical shift data, we could determine the linkage between partial helical structures and primary nucleation using our regression method.

Our regression method provides several major structures and proportion of ensemble for each structure. It is developed by focusing to fit chemical shift prediction to experimental chemical shift, which used the data from both NMR chemical shift and MD simulation with replica exchange method. It suggests the ensemble structure in a specific environment. As a result, our regression approach provides the insight of Aβ ensemble structure by bridging between NMR chemical shift and MD simulation.

Also, our method can be applied various other studies. For example, we can get structural information about major populations. We can perform further studies such as protein-protein or protein-ligand binding MD simulation using given IDP-like structures as initial structures. Some monomers or more complex protein systems are important for pharmaceutical fields. Our mathematical tool can be easily applied to other molecules including amyloid proteins, metal ions, proteins including antibody and potent inhibitors so that we can easily get the insight of molecular and thermodynamical features.

Although our study already shows some reasonable molecular features for Aβ protein, it still has some limitations and capability of improvements. The sampled structures from the regression have the potential to show specific Aβ conformation state at the time when it progresses into oligomerization. However, the results of MD simulation for Aβ monomer do not provide any intra-molecular interactions so that the chemical shift prediction does not consider the shielding effects for dimer, trimer, and oligomer state of Aβ protein. If we perform to enlarge the input ensemble pool for regression method, then we can catch out more reliable Aβ protein progressing dimerization and oligomerization. In addition, because our regression approach depends on sampling performance of MD simulation (the explored ensemble states) and accuracy of chemical shift prediction algorithm, the prediction errors can be caused. Better sampling and better chemical shift prediction would provide more reasonable ensemble structure of proteins. Moreover, there is another approach to improve the reliability of our regression approach, which is to combine previous NMR experimental data such as NOEs, *J* couplings.^14–15, 36, 57, 67^

## Conclusions

Heterogeneous and short-lived structures in ensembles are fundamental to unraveling the inherent dynamics and function as well as disease-causing aggregation of proteins. We developed the new mathematical tool to reveal compelling ensemble structures of intrinsically-disordered Aβ based on the experimental NMR chemical shift and MD simulation data. Initial Aβ structures provided possible explanations for the association and aggregation of Aβ at the atomic resolution key to understanding of AD pathogenesis. The successful application to folded proteins supports a broader exploitation of our regression approach to study ensemble structures of proteins in solution. Convincing ensemble structures with our regression will particularly contribute to gaining insights into the aggregation mechanism as well as the intermolecular interaction between amyloid proteins and potent inhibitors by supplying a reasonable template structure for the *in silico* study.

## Author contributions

W-J.Y., B.S.K, Y.-H.L., and W-K.Y. designed the research. W-J.Y., B.S.K., Y.L., and D.I. performed the research. W-J.Y., B.S.K, Y.L., D.I., J.H.K., Y.-H.L. and W-K.Y. interpreted the data. W-J.Y., B.S.K, Y.-H.L., and W-K.Y. wrote the paper with input from all authors.

## Acknowledgement

We thank the DGIST supercomputing and big data center for the allocation of dedicated supercomputing time. We are also grateful to Prof. Masahiro Hoshino (Kyoto Univ., Japan) for critical reading of the manuscript and valuable comments and to Prof. József Kardos (Eötvös Loránd Univ., Hungary) for providing ^15^N-labeld Aβ.

