## Supplemental material for "Exploring ensemble structures of Alzheimer’s amyloid β (1-42) monomer using linear regression for the MD simulation and NMR chemical shift"

#### Supplementary materials and methods

##### Materials

Sodium hydroxide and hydrochloric acid solution were purchased from Merck Millipore (Darmstadt, Germany). Isotope-unlabeled and $^{15}N$-labeled full-length Aβ42 were recombinantly expressed in *E. coli* and purified as reported previously. All the other reagents were obtained from Sigma-Aldrich (Milwaukee, WI, USA). Expression and purification of Aβ42 were performed as described previously (ref).

##### Preparation of Aβ42 sample solution

In order to remove preformed aggregates, purified Aβ42 peptides were dissolved in hexafluoroisopropanol (HFIP) and the resulting solution was aliquoted, lyophilized for 24 hours, and stored at -20 ºC. A sample solution of Aβ42 monomers was then prepared by dissolving the lyophilized peptide in 10 mM NaOH solution to a stock concentration of ca. 350 μM. The concentration of Aβ42 was determined by measuring the ultraviolet absorbance at 280 nm with the extinction coefficient of 1490 $M^{-1}$ $cm^{-1}$. Fresh Aβ42 monomer sample was diluted to a desired concentration of Aβ42 in a buffered solution (20 mM sodium phosphate, pH 7.5) for each subsequent experiment.

##### Nuclear magnetic resonance spectroscopy

Two-dimensional (2D) nuclear magnetic resonance (NMR) measurements were performed on a Bruker Avance II 800 NMR spectrometer equipped with a cryogenic probe (Bruker BioSpin, Germany). Sample of $^{15}N$-labeled Aβ42 monomer (15 μM) was prepared in 20 mM sodium phosphate buffer at pH 7.5 containing 10% $D_2O$ (volume/volume (v/v) percentage). In order to inhibit the aggregation process, the experimental temperature was maintained at 10 °C. The $^{1}H$-$^{15}N$ band-Selective Optimized-Flip-Angle Short Transient (SOFAST) heteronuclear multiple quantum coherence (HMQC) spectrum was acquired using 128 t1 experiments and 384 scans. Data processing was performed with NMRpipe[1] and Sparky 3.115.[2] The assignment of $^{15}N$-labeled Aβ42 peptide was according to the previous results.[3]

**Circular dichroism spectroscopy**

CD measurements of Aβ42 monomer (20 μM) in 20 mM sodium phosphate buffer (pH 7.5) were carried out on a JASCO J-710 spectropolarimeter (JASCO, Tokyo, Japan) using a quartz cuvette with a 0.1 cm path length. The cuvette temperature was kept to 25 °C by a water-circulating cell holder. Far-UV CD spectra between 195 and 250 nm were acquired from 16 accumulated scans using a bandwidth of 1.0 nm and an averaging time of 2 sec. CD signals were displayed in the unit of the mean residue ellipticity, [$\theta$] (deg $cm^2$ $dmol^{-1}$) after the subtraction of the solvent background. After CD measurements, BeStSel[4] was performed to determine secondary structure and fold recognition.

**Dynamic light scattering measurements**

A Zetasizer NanoZS (Malvern Panalytical Ltd., Malvern, UK) was used to assess the hydrodynamic radius of 20 μM Aβ42 monomers in 20 mm sodium phosphate buffer at pH 7.5. Aβ42 sample solution was loaded to the folded capillary zeta cell after a 5-min centrifugation at 10,000 *g*. The temperature was set to 10 °C. The average value over 10 measurements was used. Experimental data were analyzed by Zetasizer software (Malvern Panalytical Ltd., ver. 7.12).

**Molecular dynamics simulation**

***System preparation***

All systems were built using LeaP program, and all simulations were performed in AMBER20 MD simulation package.[5] The Amber ff99SBildn force field[6] was used for all simulations. The hydrogen atoms were constrained by the SHAKE algorithm.[7-8] To generate ensembles in implicit solvent model, stretched structure of Aβ42 was generated in LeaP program, ubiquitin and chymotrypsin inhibitor II (CI2) structure were obtained from Protein Data Bank (PDB) (PDB code: 3CI2[9], 1D3Z[10], respectively). The Generalized Born (GB)-Neck2[11] implicit solvent model (igb=8 in AMBER) with mbondi3[11] set was used. To use ionic strength effects, salt concentration was 150mM based on Debye-Hückel screening.[12] For Aβ in explicit solvent model, structure was obtained from PDB (PDB code: 1IYT[13]). Aβ molecules were solvated by TIP3P water model[14] with periodic boundary condition (PBC) using LeaP program in AMBER20 and those systems including 8167 water molecules were neutralized with $Na^+$ ions.

##### ***Minimization and Equilibration***

Each system was minimized with 1000 steepest descent and maximum 1000 conjugate gradient minimization steps. After minimization steps, the systems were heated during 5 ns with 2 fs time step from 20 K to each target temperature. Temperature was regulated by Langevin thermostat with 1.0 $ps^{-1}$ collision frequency.

##### ***Replica Exchange MD***

To generate ensemble structure, replica exchange molecular dynacmis (REMD)[15] simulations were carried out by PMEMD program in AMBER20.[5] Each temperature value for T-REMD simulation was generated by temperature generator for REMD-simulations.[16] For Aβ, ubiquitin and CI2 with implicit solvent model, each system which has total 10 replicas was simulated with temperature range 295 K to 544 K, 295K to 450K and 295K to 480K, respectively. The final average exchange ratios were 12.2%, 12.8%, and 10.4%, respectively. For Aβ in explicit water, total 40 replicas were simulated and their temperature was from 298 K to 417 K. The final average exchange ratio was 15.9%. During all simulations, exchanges were attempted every 10 ps and each ensemble was simulated with 1 μs. Total sampling time of implicit and explicit solvent system was 10 μs and 40 μs, respectively. Nonbonded long-range interaction was regulated by particle-mesh Ewald (PME) method[17] with 10 Å cutoff distance in explicit solvent systems.

##### ***MD trajectories analysis***

All trajectories were processed and analyzed using CPPTRAJ provided by AMBER20 package.[5] The trajectories after 200 ns were used in radius of gyration ($R_g$), contact map, and secondary structure analysis. The contact maps were calculated by distance between Cα-Cα atoms with threshold 7.5 Å. Secondary structure was analyzed by STRIDE program.[18-19] All snapshots of the trajectories were visualized using VMD.[20]

##### **Chemical shift prediction**

Prediction of chemical shift data was performed by SHIFTX2[21], SPARTA+[22] and UCBSHIFT[23]. Those algorithms predict both the backbone and side chain $^{1}H$, $^{13}C$ and $^{15}N$ chemical shifts for proteins using the PDB input files. The inputs of those algorithms were the fully equilibrated structures along the trajectories, which were total 80000 PDB inputs from 200 ns to 1000 ns for each implicit/explicit solvent MD simulation. $^{1}H$, $^{15}N$ values for each residue were selected to compare with the experiment. Incomplete $^{1}H$, $^{15}N$ data for the residues were not used. pH 7.5 condition was applied on SHIFTX2, SPARTA+ and UCBSHIFT. 283K temperature condition was applied on SHIFTX2.

### Supplementary text

#### Regression approach

##### *Evaluation of regression*

We performed multiple linear regression with predicted chemical shift data for each implicit and explicit solvent trajectory to fit the experimental chemical shift. The constraints of the multiple linear regression are: 1) the sum of coefficients is equal to 1, 2) each coefficient has a positive value and 3) the multiple linear regression has no interception. 2) & 3) constraints could be achieved by using non-negative least squares (NNLS) regression. And it acquires more modification to perform the sum of regression coefficients to be 1. Also, we tested the well-known regression method, LASSO regression, which composes both variable selection and regularization to increase the prediction accuracy and reduce the computational cost. NNLS and LASSO regressions were performed by using NumPy[24] and SciPy[25] module in Python 3.6. Mathematically, the notations of NNLS regression and LASSO regression are similar as below:

$$NNLS : \underset{\boldsymbol{w}}{argmin} \; \left\|\boldsymbol{y}^T - X\boldsymbol{w}^T\right\|_2 \text{ for } w_i \geq 0, \forall \text{i} \tag{S1}$$

$$\text{LASSO} : \underset{\boldsymbol{w}}{argmin} \; \left\{ {}^{1}\!/_{2n_{res}} \left\|\boldsymbol{y}^T - X\boldsymbol{w}^T\right\|_2^2 + \alpha\left\|\boldsymbol{w}\right\|_1 \right\} \tag{S2}$$

$\boldsymbol{y}$ is an experimental chemical shift, which is a $2n_{res}$ —dimensional vector (combined $^1$H and $^{15}$N). $n_{res}$ is the number of residues for protein in chemical shift prediction. $X$ is input chemical shift prediction dataset, which is $2n_{res} \times$ N matrix. N is the number of structures. $w_i$ is each coefficient of the regression.

If the hyper-parameter, α, sets a small positive value, then the optimal solves of LASSO converges to the result of NNLS regression. For our interpretation of the convergence of LASSO regressions, NNLS regression is more accurate in our case when we restrained all coefficients to be positive and the sum of regression coefficients to be 1. Under table (which resulted from scaled dataset) supports our argument (α=0.05, positive=True in LASSO).

| | | UCBSHIFT | | | SHIFTX2 | | |
|---|---|---|---|---|---|---|---|
| | | $Score_{total}$ | $Score_H$ | $Score_N$ | $Score_{total}$ | $Score_H$ | $Score_N$ |
| GB(Implicit) | NNLS | **.9763** | **.9901** | **.9541** | .9499 | .9812 | .9069 |
| | LASSO | .9759 | .9896 | .9536 | .9495 | .9796 | .9077 |

***Regression scoring***

The score of the multiple linear regression is based on the coefficient of determination, denoted $R^2$, which represents the proportion of the variance about the chemical shift for each residue of Aβ monomer. The score of each atom in chemical shift ($^1$H and $^{15}$N) is separately calculated in this scheme. The detailed scoring method is as follows:

$$RSS = \sum_{i=1}^{n} (y_i - \hat{y_i})^2, \qquad TSS = \sum_{i=1}^{n} (y_i - \bar{y})^2$$

$$R^2 = 1 - \frac{RSS}{TSS} \tag{S1}$$

$y$ is the reference chemical shift and $\bar{y}$ is a its mean value. $\hat{y}$ is the predicted chemical shift. RSS and TSS is the residual sum of square and total sum of square, respectively.

And, to quantitatively compare for each case, we define a total score combining the scores of all atoms. More precisely the equation of the total score:

(S2)

In this scoring method, if the model is exactly matched to a true reference, then RSS goes on 0 so that the $R^2$ score becomes 0. Also, this score can be negative if the model shows a worse prediction. Mathematically, it is possible if the variance of chemical shift prediction is much larger than the variance of reference, which means the averaged-out, denoted non-weighted coefficient, model is actually wrong.

***Dataset scaling***

We preformed scaling process for the chemical shift data because of the scale difference between $^{1}$H and $^{15}$N values. Because of the scale difference, normal NNLS regression results showed that $^{15}$N values (which have much bigger scale than $^{1}$H values) are more well-fitted before scaling process. Therefore, this scaling process gives us more increased accuracy of regression. Graphics were plotted by matplotlib[26].

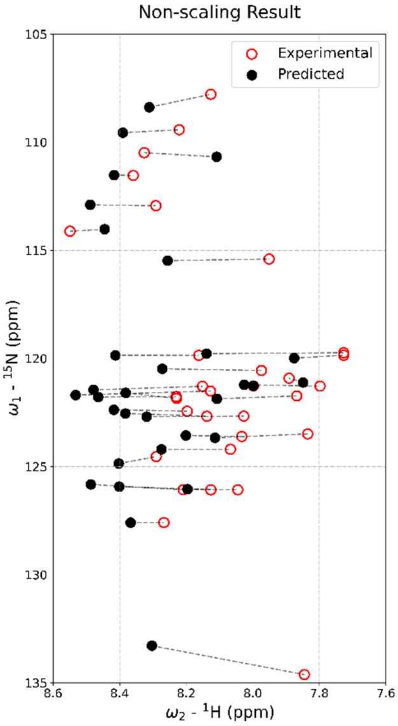

**Non-scaling result.** The result of non-negative least squares regression using non-scaled input. All data points only adopt to the $^{1}$H axis because the average value of $^{1}$H axis is much smaller than $^{15}$N average

The point of regardless is that the result of regression for scaling dataset can be ensured an accurate prediction for non-scaling original dataset. In addition, we should check that it is possible to directly apply the coefficients of regression for scaled dataset to original dataset without any additional modification. We could prove it by showing the existence of inverse scaling function. Scaling function $f$ is as follows:

$$\boldsymbol{y} = \widehat{\boldsymbol{y}} + \epsilon = w_1 X_1 + w_2 X_2 + \cdots + w_n X_n + \epsilon, \tag{S3}$$

$$\hat{\boldsymbol{y}}^T = \begin{bmatrix} \hat{y}^{1H} \\ \hat{y}^{1N} \\ \hat{y}^{2H} \\ \hat{y}^{2N} \\ \vdots \\ \hat{y}^{nH} \\ \hat{y}^{nH} \end{bmatrix}, \quad X_k^T = \begin{bmatrix} x_k^{1H} \\ x_k^{1N} \\ x_k^{2H} \\ x_k^{2N} \\ \vdots \\ x_k^{m\,H} \\ x_k^{m\,N} \end{bmatrix}$$

$$f(X_k^T) = \begin{bmatrix} w_H \cdot \left(x_k^{1H} - x_{avg}^{H}\right) \\ w_N \cdot \left(x_k^{1N} - x_{avg}^{N}\right) \\ w_H \cdot \left(x_k^{2H} - x_{avg}^{H}\right) \\ w_N \cdot \left(x_k^{2N} - x_{avg}^{N}\right) \\ \vdots \\ w_H \cdot \left(x_k^{m\,H} - x_{avg}^{H}\right) \\ w_N \cdot \left(x_k^{m\,N} - x_{avg}^{N}\right) \end{bmatrix} \tag{S4}$$

$$x_{avg}^{H} = \frac{1}{nm}\sum_{k=1}^{n}\sum_{l=1}^{m} x_k^{m\,H}, \qquad x_{avg}^{N} = \frac{1}{nm}\sum_{k=1}^{n}\sum_{l=1}^{m} x_k^{m\,N} \tag{S5}$$

$$w_{\{H,N\}} = \{1, \ \frac{1}{30}\ \} \tag{S6}$$

$\hat{\boldsymbol{y}}$ is predicted chemical shift among the ensemble of protein. $\epsilon$ is error value. Empirical scaling factors, $w_{\{H,N\}}$, represent the range of chemical shift in each $^{1}$H, $^{15}$N axis. Regularization factor was used as $x_{avg}^{\{H,N\}}$. We can simplify above expression as follows:

$$\hat{\boldsymbol{y}}_f = \left(\sum_{k=1}^{n} w_k f(X_k)\right)^T = \begin{bmatrix} w_H \\ w_N \\ \vdots \\ w_H \\ w_N \end{bmatrix} \odot \sum_{k=1}^{n} w_k \left( X_k^T - \begin{bmatrix} x_{avg}^{H} \\ x_{avg}^{N} \\ \vdots \\ x_{avg}^{H} \\ x_{avg}^{N} \end{bmatrix} \right) = (f \circ \hat{\boldsymbol{y}})_i = f(\hat{\boldsymbol{y}}) \tag{S7}$$

$\odot$ is element-wise product, called Hadamard product. It is as same as multiplying a vector by a diagonal matrix (i.e., $\mathrm{diag}(w_H, w_N, \dots, w_H, w_N)$). Definitely, it has an inverse matrix. Thus, it satisfies:

$$\begin{bmatrix} 1/w_H \\ 1/w_N \\ \vdots \\ 1/w_H \\ 1/w_N \end{bmatrix} \odot \left(\sum_{k=1}^{n} w_k f(X_k)\right)^T + \sum_{k=1}^{n} w_k \begin{bmatrix} x_{avg}^H \\ x_{avg}^N \\ \vdots \\ x_{avg}^H \\ x_{avg}^N \end{bmatrix} := \left(\sum_{k=1}^{n} w_k f^{-1} \circ f(X_k)\right)^T = \left(\sum_{k=1}^{n} w_k X_k\right)^T \quad \text{(S8)}$$

$$\Leftrightarrow \left(f^{-1} \circ \hat{\boldsymbol{y}}_f\right)_i = f^{-1}((f \circ \hat{\boldsymbol{y}})_i) = \hat{\boldsymbol{y}} \quad \text{(S9)}$$

In result, the inverse of scaling function, $f^{-1}$, already exists so that we can directly apply the coefficients of regression for scaled dataset to original dataset.

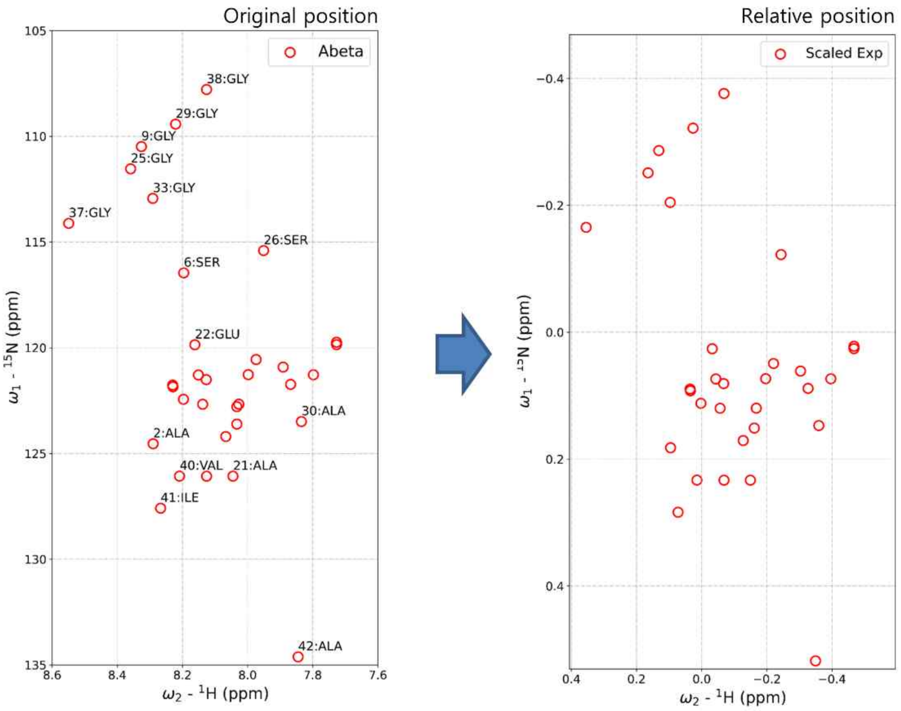

**Scaling process result**. After scaling process, $^{1}H$ values and $^{15}N$ values have similar scales so that they are comparable. Because the inverse of scaling function $\boldsymbol{f^{-1}}$ exists, we can regenerate the original position from scaling function space

***Hyperparameter fine-tuning for NMR chemical shift***

Our regression approach is highly sensitive to the true reference data because NNLS regression approach performs to minimize the residual sum of square (RSS) for reference data. However, $^{1}H$ NMR chemical shift is influenced by the environment such as temperature, pH, pressure, magnet susceptibility of machine, …, etc. To solve this problem, we introduced a specific hyperparameter $\delta(\boldsymbol{\Theta})$.

$$\boldsymbol{\Theta} = (Temp., pH, pressure, \ldots) = (\theta_1, \theta_2, \theta_3, \cdots, \theta_d) \subset \mathbb{R}^d \qquad \text{(S10)}$$

Our purpose is to project the chemical shift data onto a regularized space.

$$\boldsymbol{y}\left(\boldsymbol{\Theta} = \overrightarrow{\Theta_?}\right) = \sum_{k=1}^{n} w_k \left(X_k + g\big(\delta(\boldsymbol{\Theta})\big)\right) + \varepsilon \tag{S11}$$

$$g\big(\delta(\boldsymbol{\Theta})\big) = \boldsymbol{diag}\ (1^H, 0, 1^H, 0, \cdots, 1^H, 0) \cdot J_{1,2n_{res}} \cdot \delta(\boldsymbol{\Theta}) \tag{S12}$$

$g$ is a function that allows $\delta(\boldsymbol{\Theta})$ to be applied only to the $^{1}$H axis. $\overrightarrow{\Theta_?}$ is an arbitrary space on which the environmental effects can be ignored. Furthermore, this optimization process is applied after performing scaling function.

$$\boldsymbol{y}_f(\boldsymbol{\Theta} = \boldsymbol{0}) = \sum_{k=1}^{n} w_k \cdot \left\{f(X_k) + f \circ g\big(\delta(\boldsymbol{\Theta})\big)\right\} + \varepsilon_f \tag{S13}$$

We preformed this constrained optimization for $\delta(\boldsymbol{\Theta})$ by using a parallelized version of limited memory BFGS (L-BFGS) algorithm[27] which is an optimization algorithm in the family of quasi-Newton methods. We set the gradient function as the score function to minimize itself, which is defined as RSS/TSS. RSS and TSS were calculated by the reference and each chemical shift prediction.

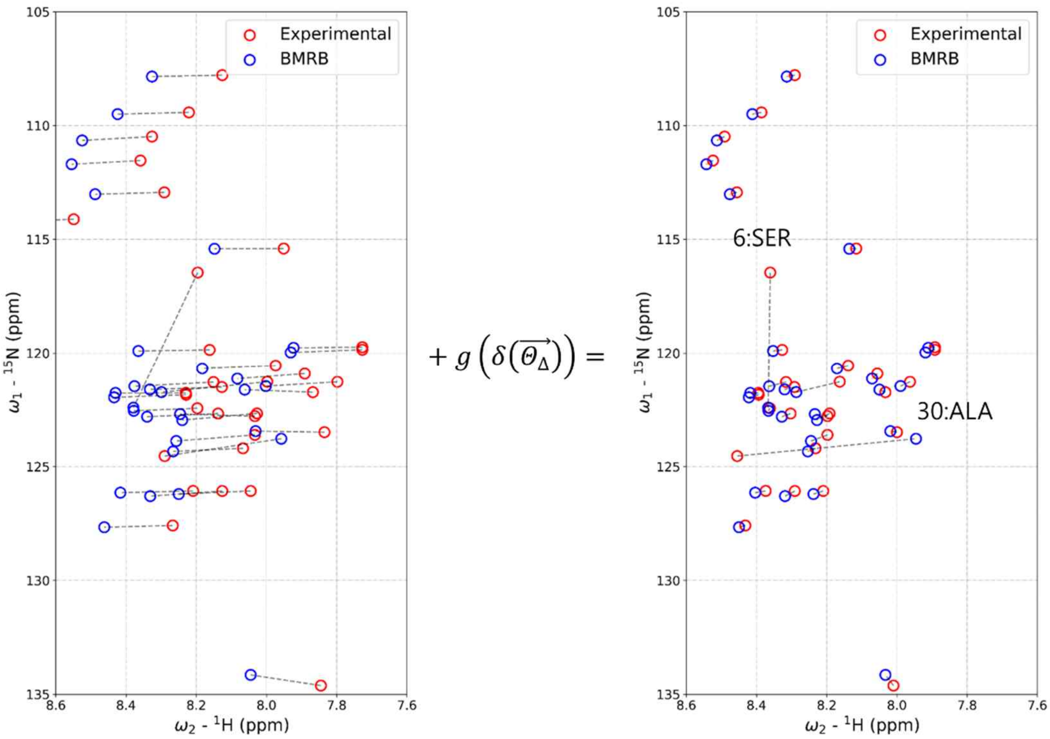

$\boldsymbol{\delta(\Theta)}$ **optimization result.** before (*left*) and after (*right*). $\boldsymbol{\delta(\Theta)}$ optimization preformed to correct the references from different sources and maximized the score function between each references and chemical shift prediction results. It allows to give consistent results for a same protein target. $\boldsymbol{\delta(\overrightarrow{\Theta_{\Delta_{Exp}}})}$ = -0.1648042, $\boldsymbol{\delta(\overrightarrow{\Theta_{\Delta_{BMRB}}})}$ = 0.0123154.

##### *Normalization of regression coefficients*

The sum of regression coefficients is not usually equal to 1. Also, a simple normalization method of regression coefficients by the sum of coefficients is not appropriate because it does not guarantee that the data will not be altered and the error is not enough small. To solve this problem, we suggested a realizable procedure. First, we selected non-negligible features of the regression based on following criteria:

$$A = \{a_i\}_{i=1}^{N} \text{ such that } a_i \geq \varepsilon \cdot a_m,\ a_m = \max_{a \in A} A \tag{S14}$$

$A$ is a set of significant coefficients. $\varepsilon$ is small positive value ($10^{-5}$) as threshold of selection. $a_m$ is the maximum element of the set $A$, which is equal to the maximum element of whole coefficient set.

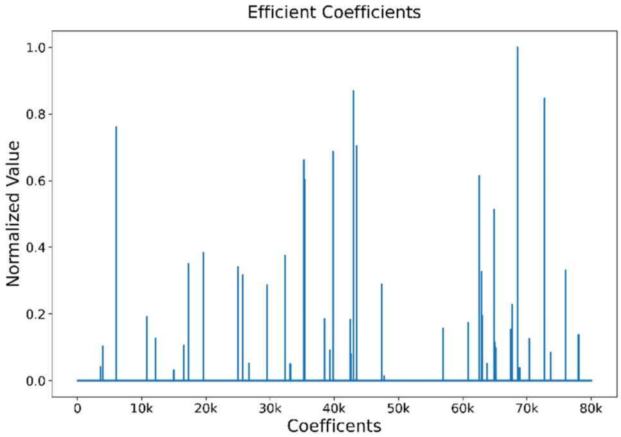

**The example distribution of regression coefficients**. Almost coefficients converge to 0 excluding some significant coefficients. Approximately, 20~50 coefficients can be efficiently selected by our policy.

After selection, sampled coefficients can compose to original chemical shift prediction. Also, this process resulted in dimensional reduction so that it allows to perform an additional constrained optimization process. The constraint, sum of coefficients should be 1, is achieved through the minimization process using SLSQP algorithm in SciPy optimization module.

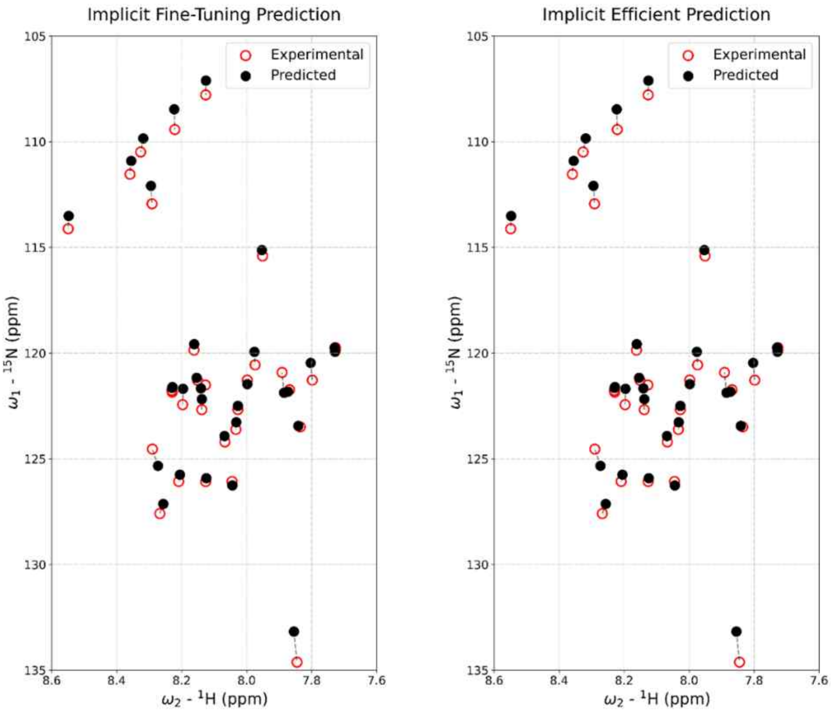

**Efficiently mimicked prediction data.** before (*left*) and after (*right*). After significant data sampling with suggested policy, we can eliminate meaningless data. This dimensional reduction method allows to perform an additional constrained minimization process.

***Careful precautions***

In general, the sum of coefficients is more than 1. If you just use a method of normalization with the sum of coefficients, then the predicted result is not stable which means the error of prediction is not enough small.

From Eq. S3, we can understand the meaning of error.

$$\hat{\boldsymbol{y}} = \sum_{i=1}^{n} w_i \left( \frac{1}{\sum_{k=1}^{n} w_k} + 1 - \frac{1}{\sum_{k=1}^{n} w_k} \right) X_i \tag{S15}$$

$$\Leftrightarrow \hat{\boldsymbol{y}} = \sum_{i=1}^{n} \frac{w_i}{\sum_{k=1}^{n} w_k} X_i + \left( 1 - \frac{1}{\sum_{k=1}^{n} w_k} \right) \sum_{i=1}^{n} w_i X_i \tag{S16}$$

Second term of Eq. S16 is the normalization error we discussed. It depends on $\hat{\boldsymbol{y}}$ because $1 - \frac{1}{\sum_{k=1}^{n} w_k}$ is a constant value. Thus, the simple normalization approach causes reference-dependent error.

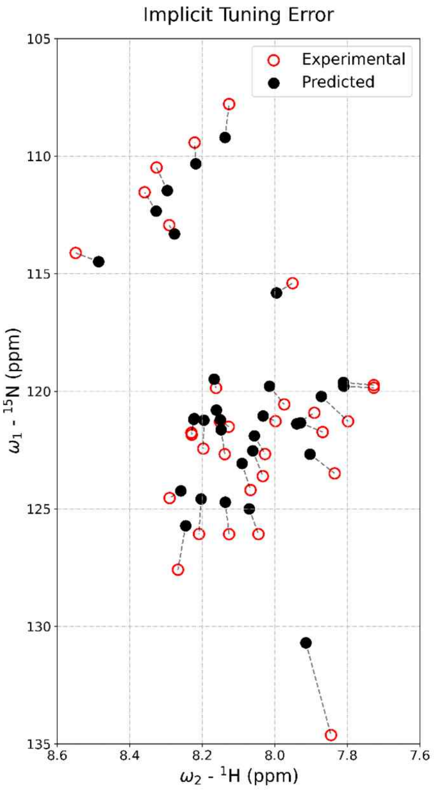

**Normalization error**. If we achieve the sum of coefficients equal to 1, we could interpret each coefficient as the state probability or the proportion of each state. But the simple ratio approach which normalizes each coefficient by the sum of coefficients value shows non-negligible error.

#### Supplementary figures

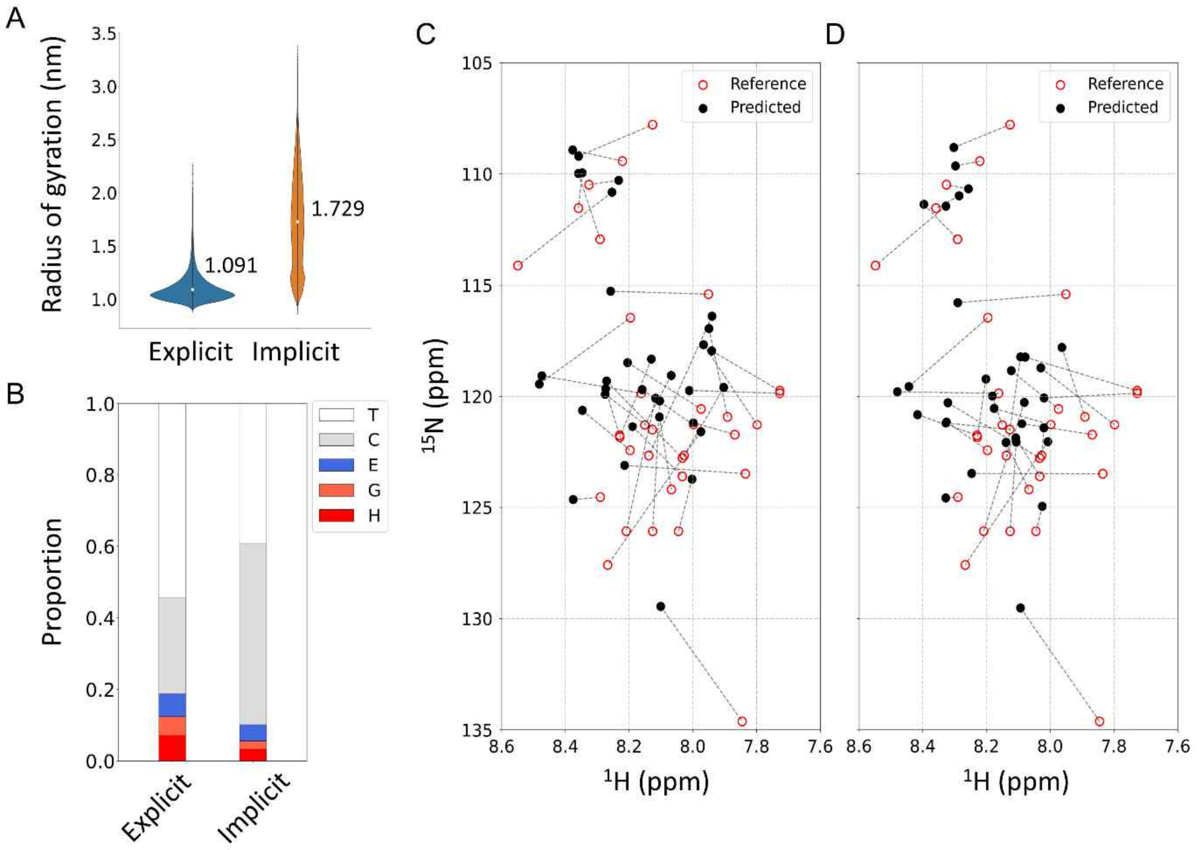

**Figure S1. structural properties of Aβ42 monomers in ensembles on both solvent models.** (A, B) The radius of gyration ($R_g$) (A) and proportion of the secondary structure (B). (A) The average values are plotted in both solvent model (*white dot*). (B) The average secondary proportions, including α-helix, $3_{10}$-helix, extended beta, coil, and turn, are represented in main text. H; α-helix, G; $3_{10}$-helix, E; β-sheet, C; coil, T; turn. (C, D) The average of predicted chemical shift for whole ensemble structures using UCBSHIFT in explicit (C) and implicit solvent model (D).

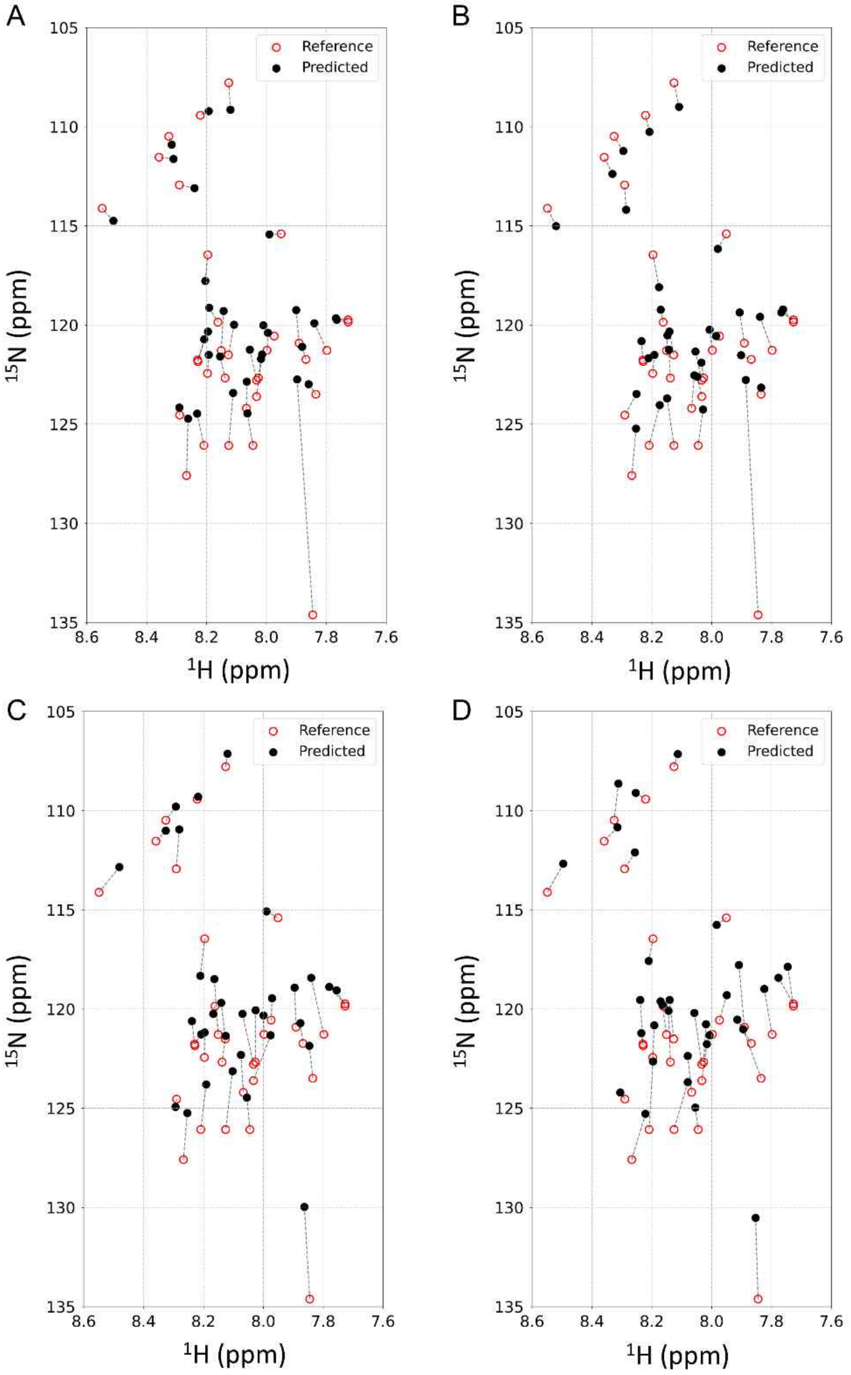

**Figure S2. Comparison between predicted and experimentally-obtained NMR chemical shifts of Aβ42.** (A-D) Weighted average of chemical shift prediction using the regression method (black) and chemical shifts obtained using HMQC measurements (red) are shown. NMR chemical shifts were obtained using SPARTA+ (A and B) and SHIFTX2 (C and D) in the explicit (A and C) and implicit solvent models (B and D).

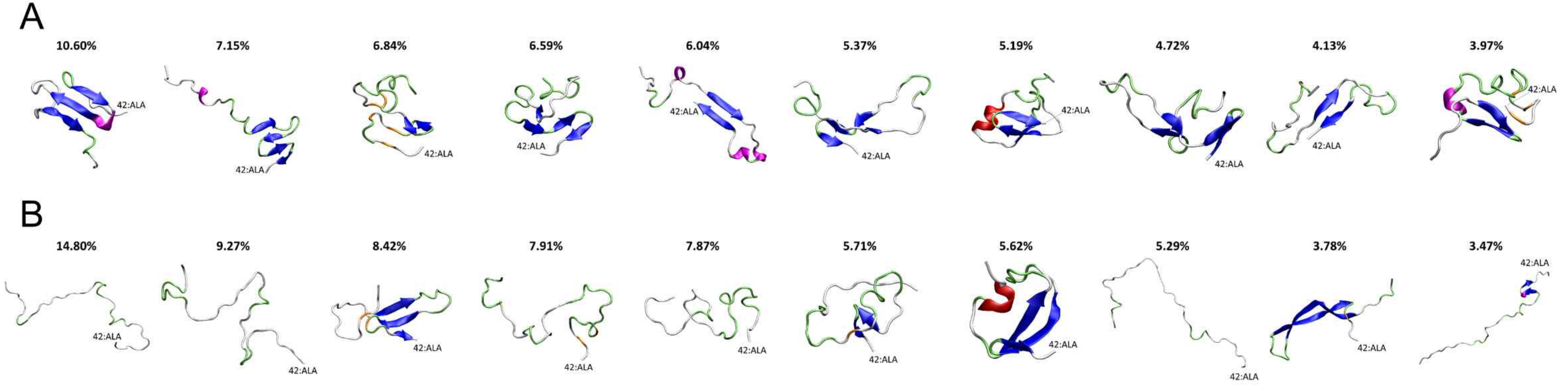

**Figure S3. Visualization of Aβ42 monomers calculated using experimental NMR chemical shifts and UCBSHIFT.** Ten representative structures obtained with the explicit (A) and implicit solvent models (B) are shown. The secondary structures are colored as follows: α-helix (red), β-sheet (blue), $3_{10}$-helix (magenta), β-bridge (orange), turn (lime), and coil (white).

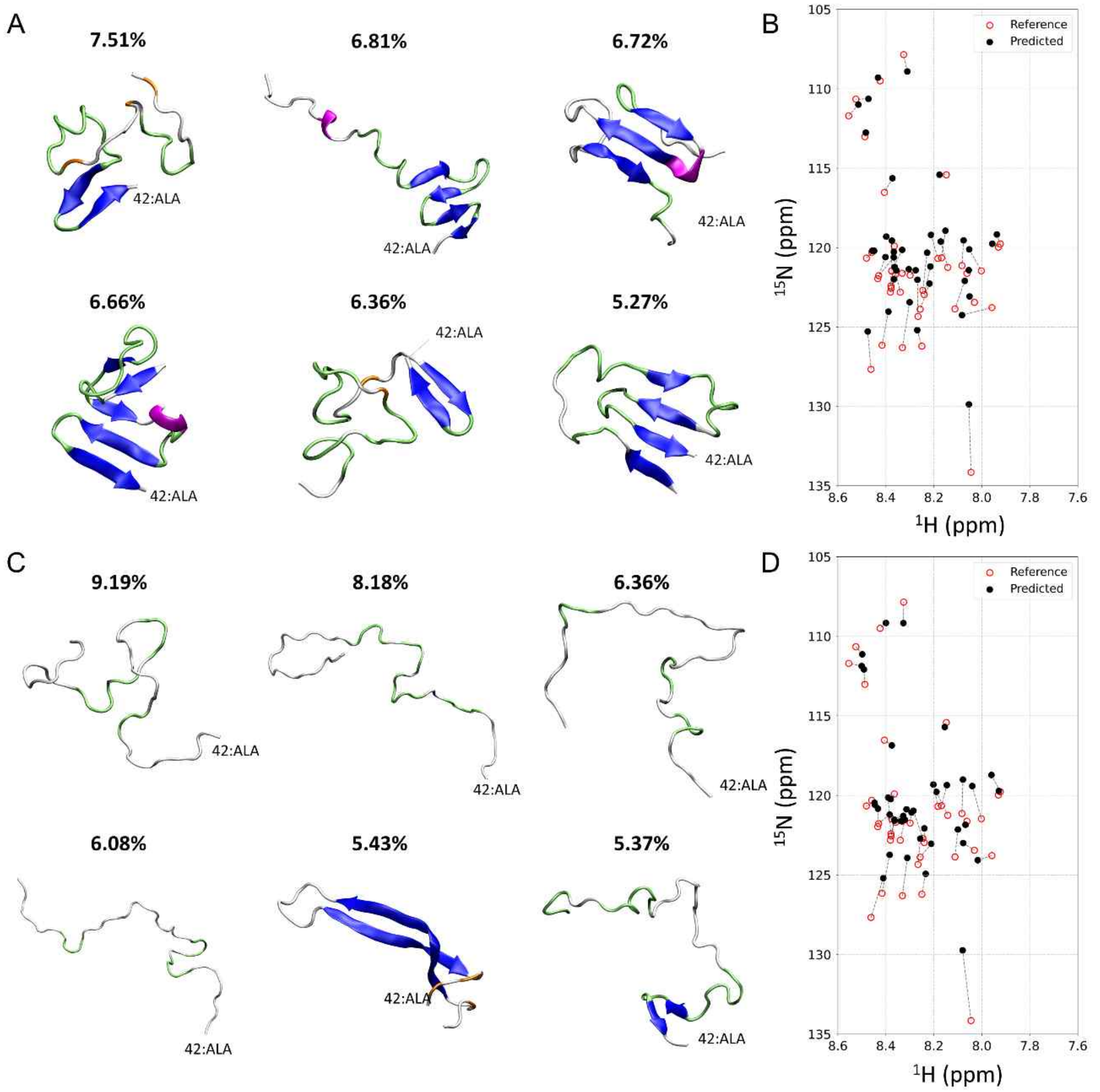

**Figure S4. The major structures of Aβ42 monomers in the ensemble for published chemical shift.** (A, C) Representative Aβ42 structures with higher populations obtained using the published NMR chemical shift (BMRB entry 25218) and UCBSHIFT in the explicit (A) and implicit solvent model (C). Secondary structures are colored as follows: α-helix (red), β-sheet (blue), $3_{10}$-helix (magenta), β-bridge (orange), turn (lime), and coil (white). The weighted average of chemical shift data by the proportion in the explicit (B) and implicit solvent model (D) is superimposed with experimentally-determined chemical shifts.

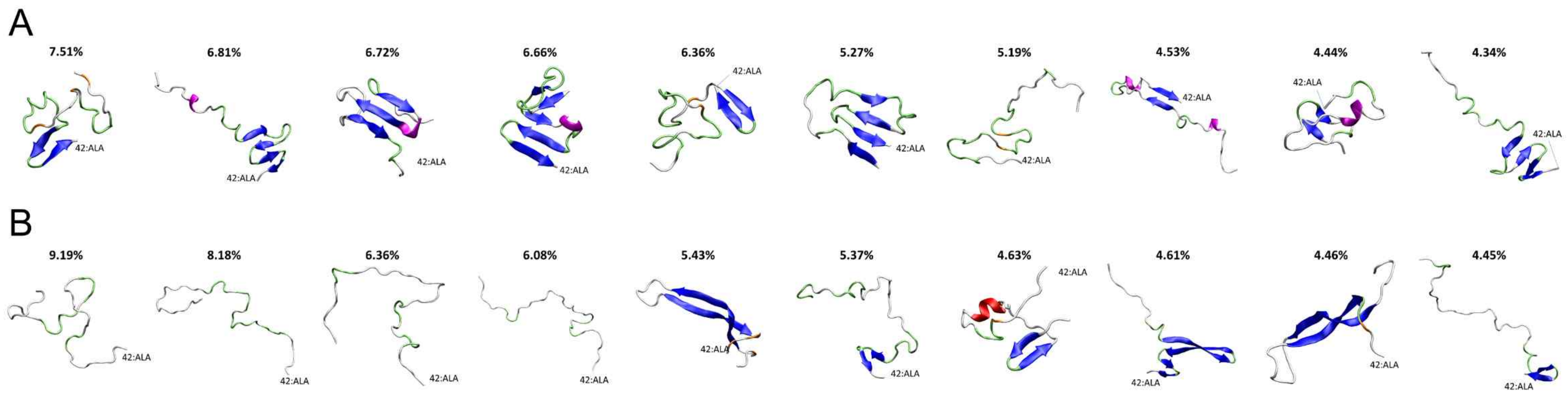

**Figure S5. Visualization of Aβ42 monomers calculated using published chemical shift and UCBSHIFT.** Ten representative structures obtained with the explicit (A) and implicit solvent models (B) are shown. Reference chemical shift is from BMRB database (BMRB entry 25218). The secondary structures are colored as follows: α-helix (red), β-sheet (blue), $3_{10}$-helix (magenta), β-bridge (orange), turn (lime), and coil (white).

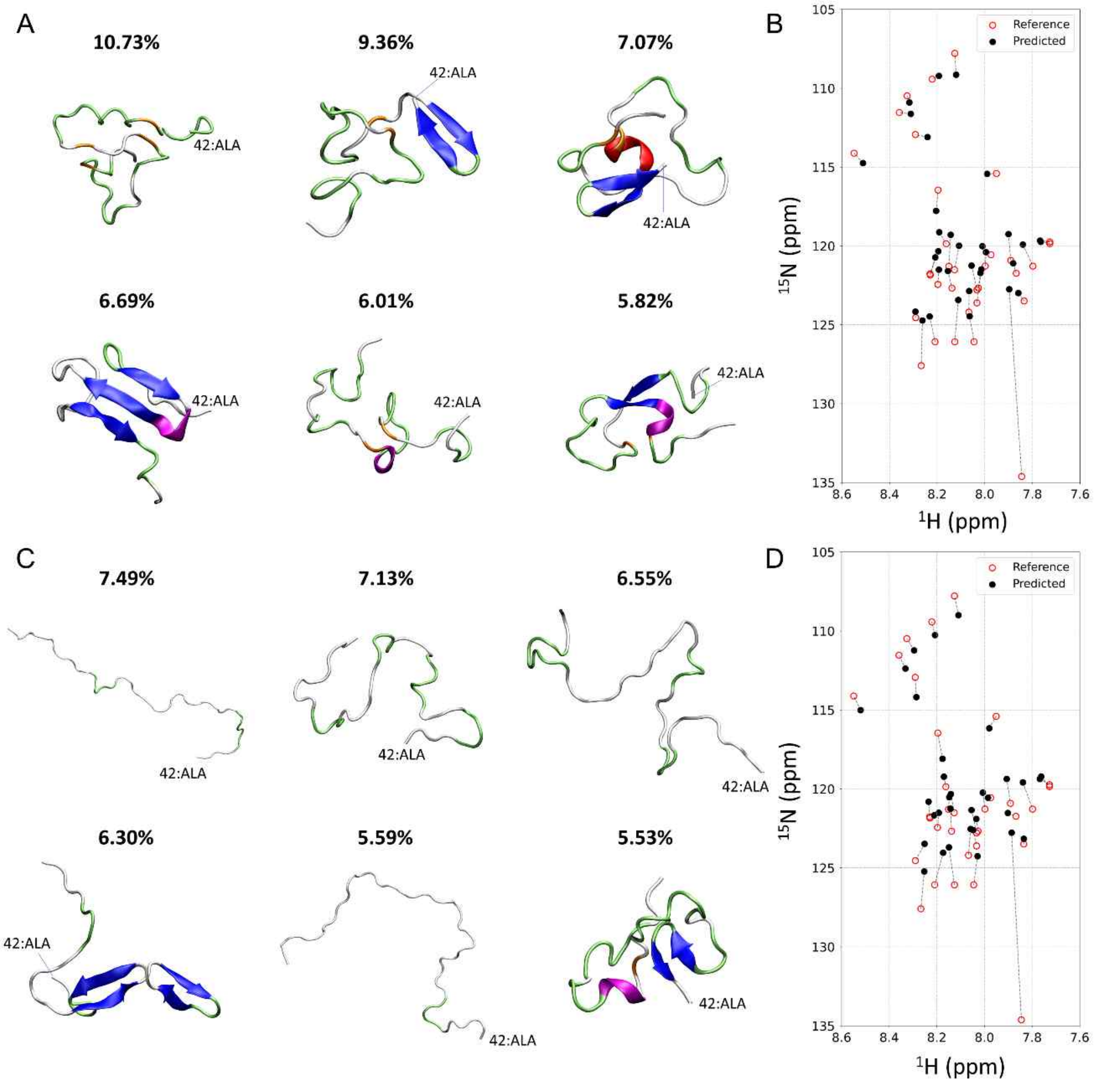

**Figure S6. The major structures of Aβ42 monomers for SPARTA+.** (A, C) Representative Aβ42 structures with higher populations obtained using the experimental NMR chemical shift and SPARTA+ in the explicit (A) and implicit solvent model (C). Secondary structures are colored as follows: α-helix (red), β-sheet (blue), $3_{10}$-helix (magenta), β-bridge (orange), turn (lime), and coil (white). The weighted average of chemical shift data by the proportion in the explicit (B) and implicit solvent model (D) is superimposed with experimentally-determined chemical shifts.

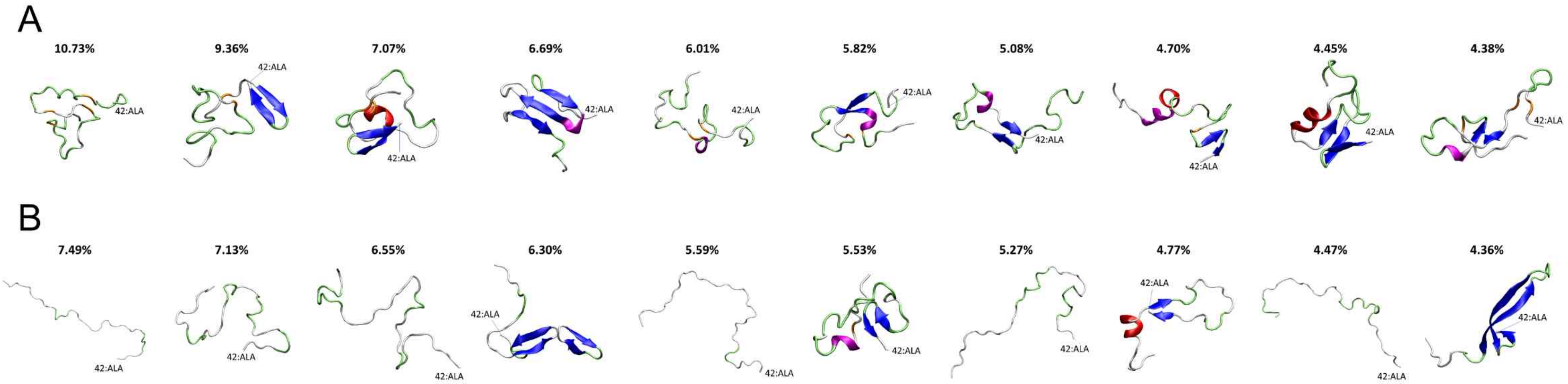

Figure S7. **Visualization of Aβ42 monomers calculated using experimental NMR chemical shifts and SPARTA+.** Ten representative structures obtained with the explicit (A) and implicit solvent models (B) are shown. The secondary structures are colored as follows: α-helix (red), β-sheet (blue), $3_{10}$-helix (magenta), β-bridge (orange), turn (lime), and coil (white).

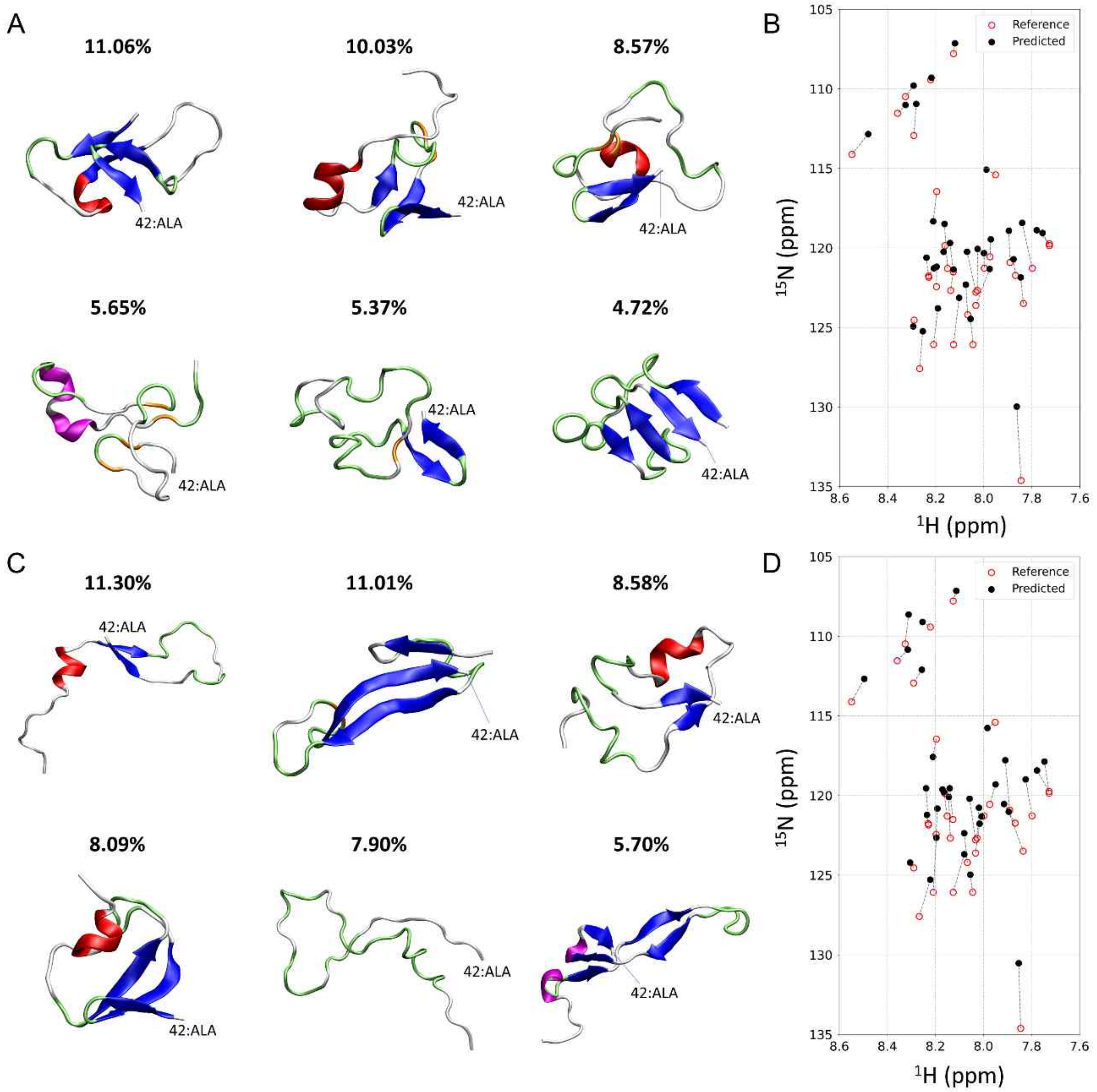

**Figure S8. The major structures of Aβ42 monomers for SHIFTX2.** (A, C) Representative Aβ42 structures with higher populations obtained using the experimental NMR chemical shift and SHIFTX2 in the explicit (A) and implicit solvent model (C). Secondary structures are colored as follows: α-helix (red), β-sheet (blue), $3_{10}$-helix (magenta), β-bridge (orange), turn (lime), and coil (white). The weighted average of chemical shift data by the proportion in the explicit (B) and implicit solvent model (D) is superimposed with experimentally-determined chemical shifts.

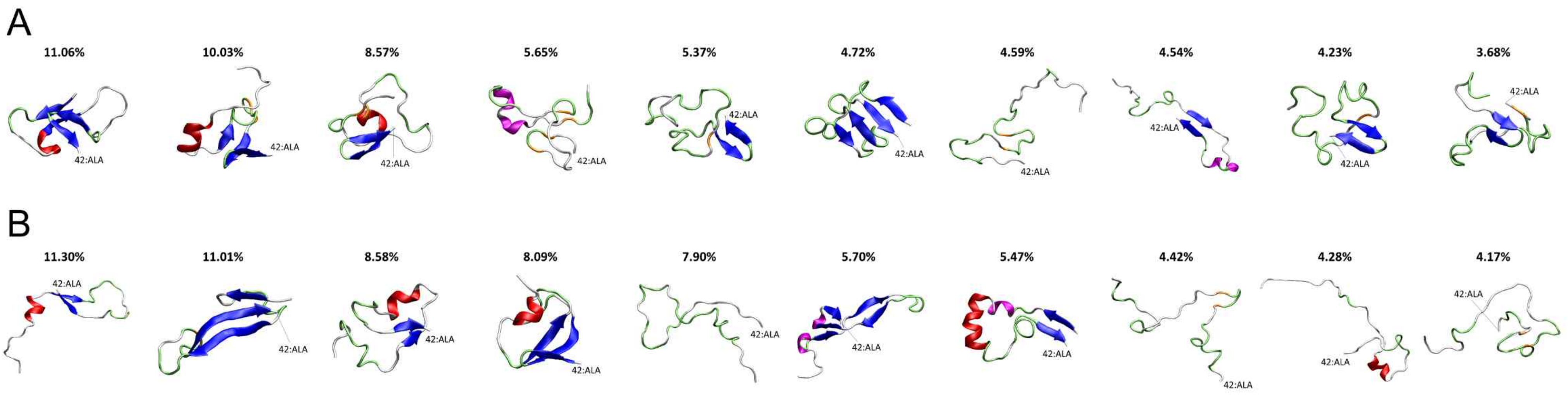

Figure S9. **Visualization of Aβ42 monomers calculated using experimental NMR chemical shifts and SHIFTX2.** Ten representative structures obtained with the explicit (A) and implicit solvent models (B) are shown. The secondary structures are colored as follows: α-helix (red), β-sheet (blue), $3_{10}$-helix (magenta), β-bridge (orange), turn (lime), and coil (white).

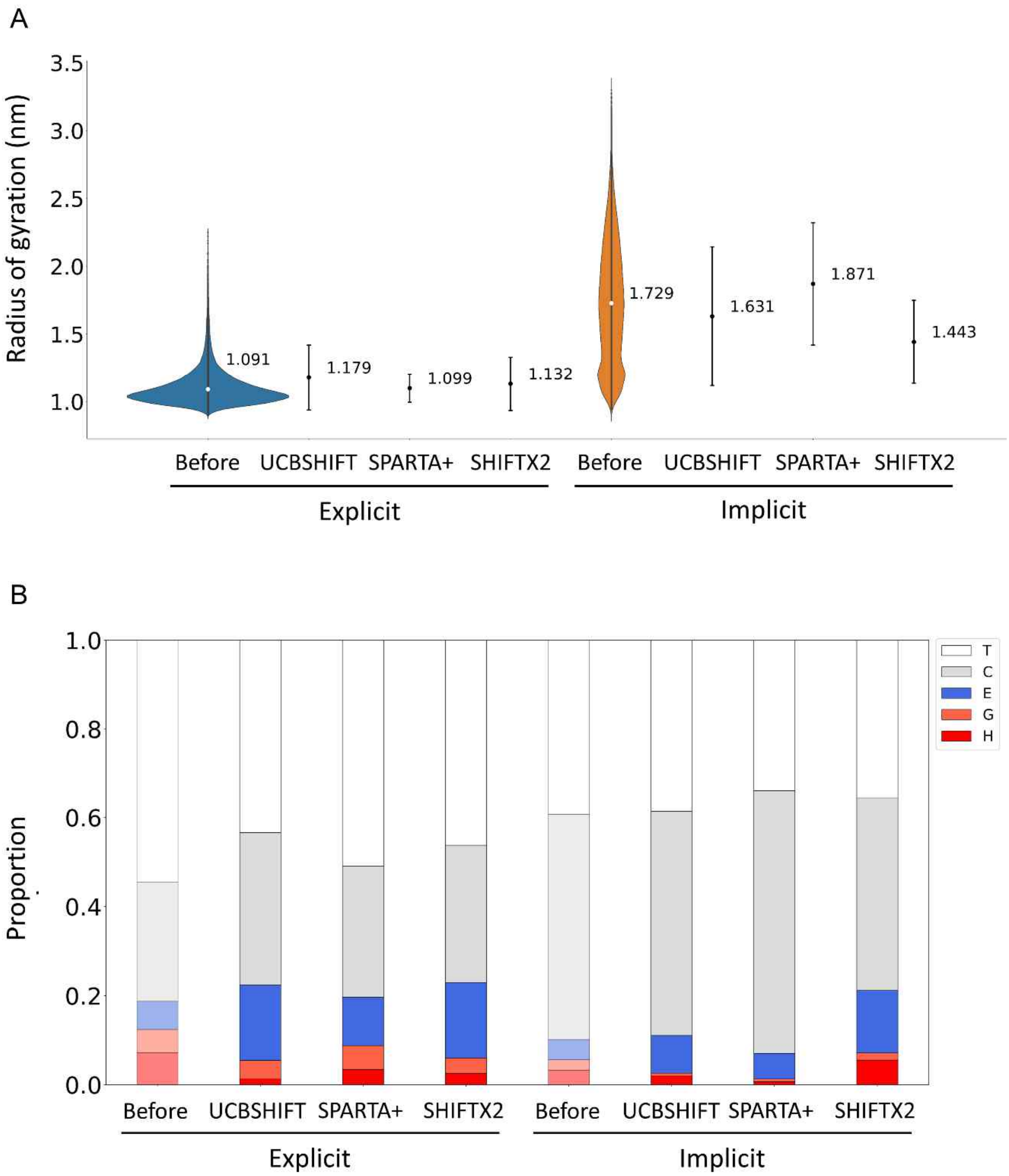

**Figure S10. Effects of linear regression on structural properties of Aβ42 monomers in ensembles for all prediction algorithms.** (A, B) The radius of gyration ($R_g$) (A) and proportion of the secondary structure (B) of Aβ42 before (Explicit and Implicit) and after multiple linear regression. Error bars in A indicate the standard deviation. The type of the secondary structure is shown with the single letter and color as follows: H, α-helix (red); G, $3_{10}$-helix (orange); E, β-sheet (blue); C, coil (gray); and T, turn (white).

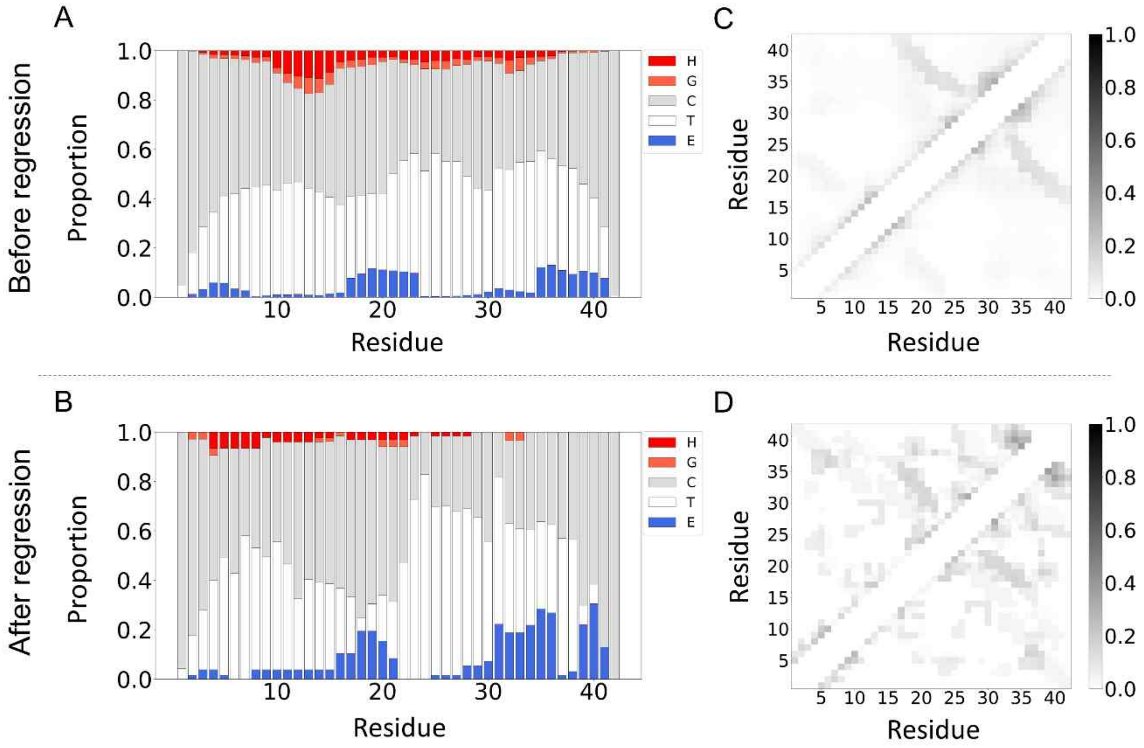

**Figure S11. Linear regression-dependent changes in the secondary structure and intramolecular contact of Aβ42.** Proportion of the secondary structure (A and B) and contact map (C and D) of Aβ42 averaged in the ensemble before (A and C) and after the regression approach (B and D) in the implicit solvent model. The type of the secondary structure in A and B is shown with the single letter and color as follows: H, α-helix (red); G, $3_{10}$-helix (orange); E, β-sheet (blue); C, coil (gray); and T, turn (white). Degree of the contact between residues of Aβ42 in C and D is scaled from 0.0 to 1.0 which is shown by the gradation (right).

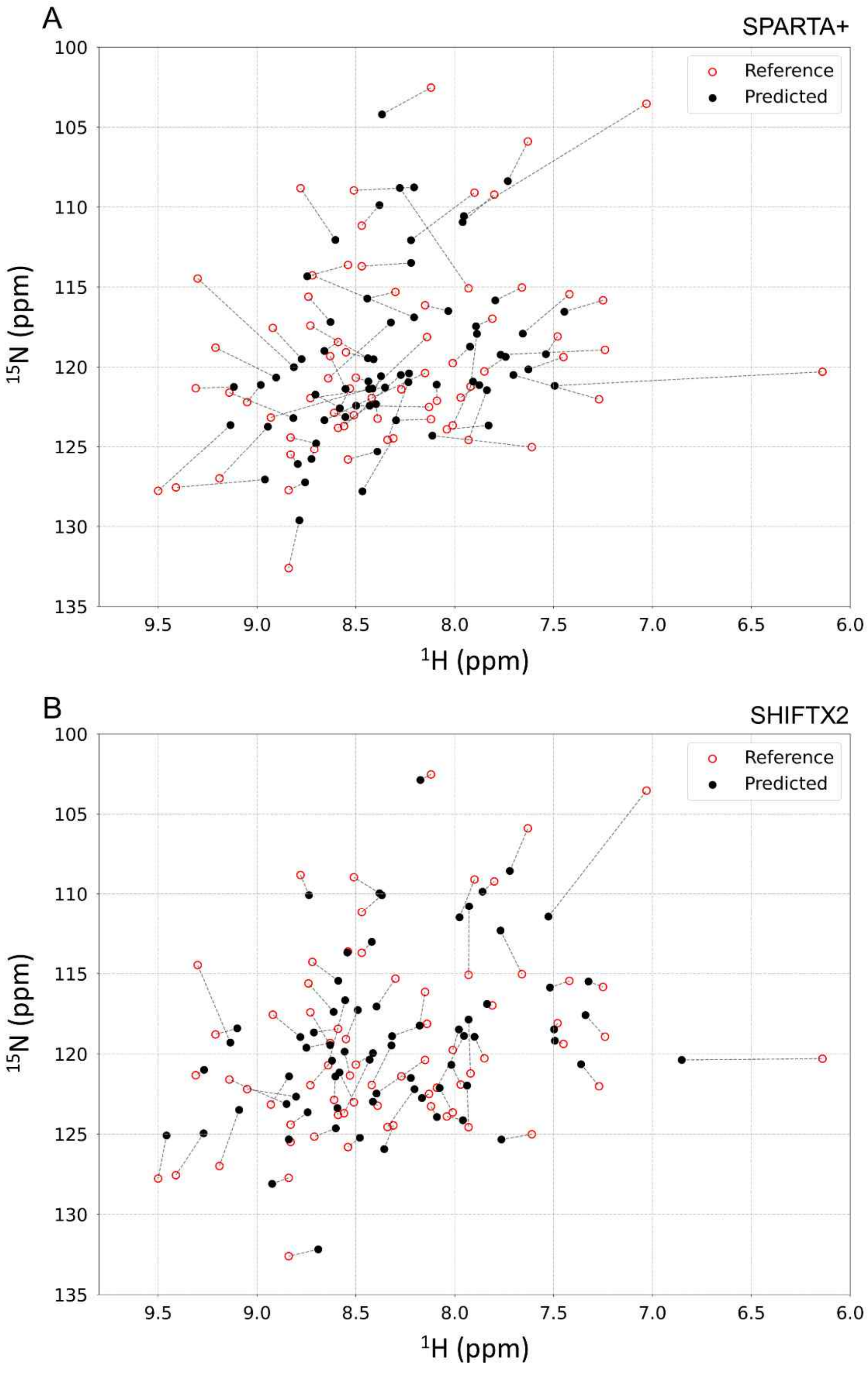

**Figure S1122. Comparison between predicted and experimentally-obtained NMR chemical shifts of ubiquitin.** (A and B) Weighted average of chemical shift prediction using the regression method (black) and chemical shifts obtained using HSQC measurements (red) are shown. NMR chemical shifts were obtained using SPARTA+ (A) and SHIFTX2 (B).

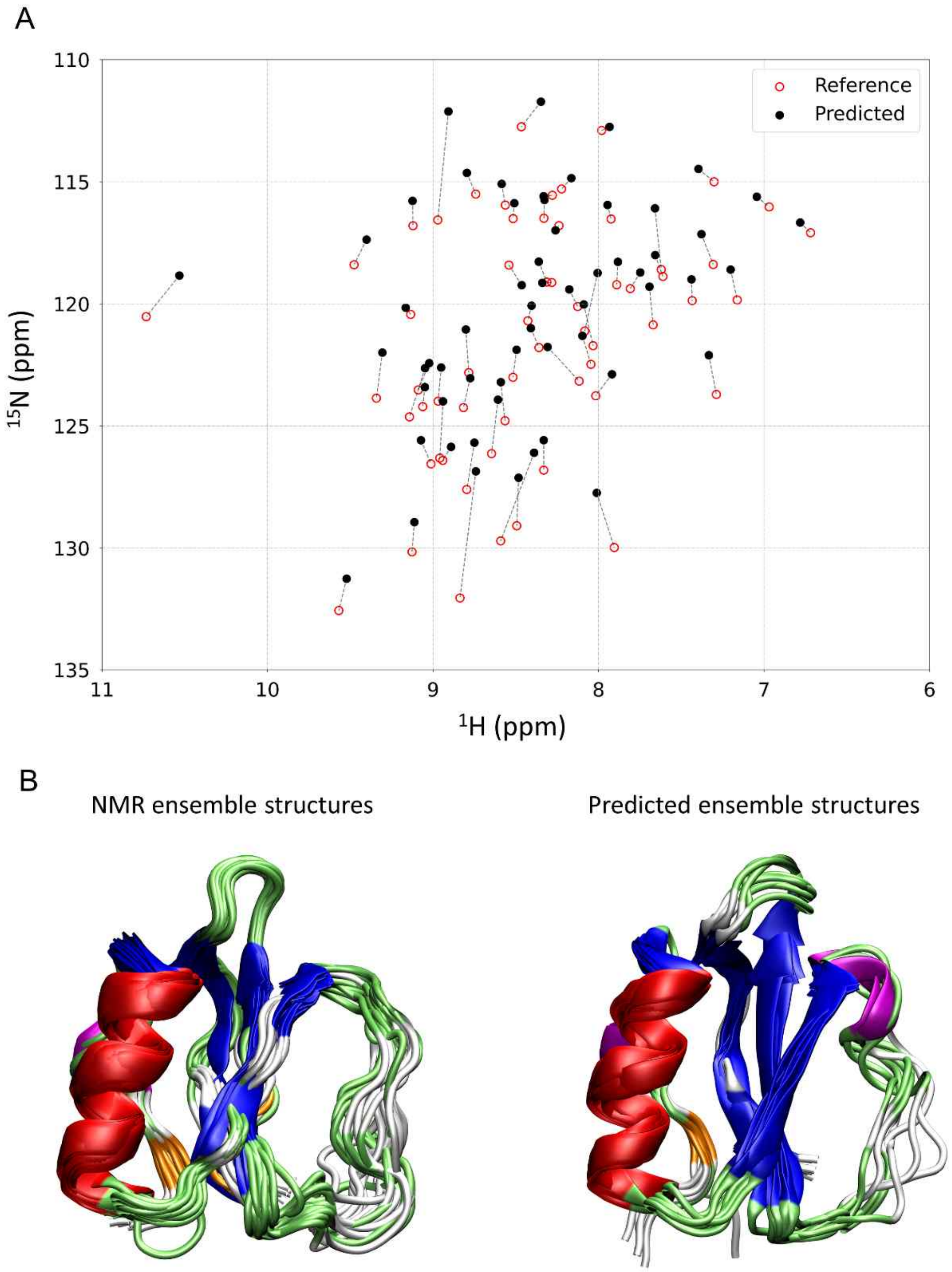

**Figure S13. Multiple linear regression approach for CI2.** (A) Comparison of chemical shifts calculated using linear regression (red) with those directly obtained using NMR spectroscopy (black) (BMRB Entry 4974. (B) Comparison between NMR-based and predicted ensemble structures of folded proteins. Three dimensional structures of CI2 (PDB code: 3CI2, BMRB Entry 4974) are shown with color codes for the secondary structures: α-helix (red), β-sheet (blue), $3_{10}$-helix (magenta), β-bridge (orange), turn (lime), and coil (white). Structures of both proteins were determined using solution NMR spectroscopy (left) and the linear regression method (right).

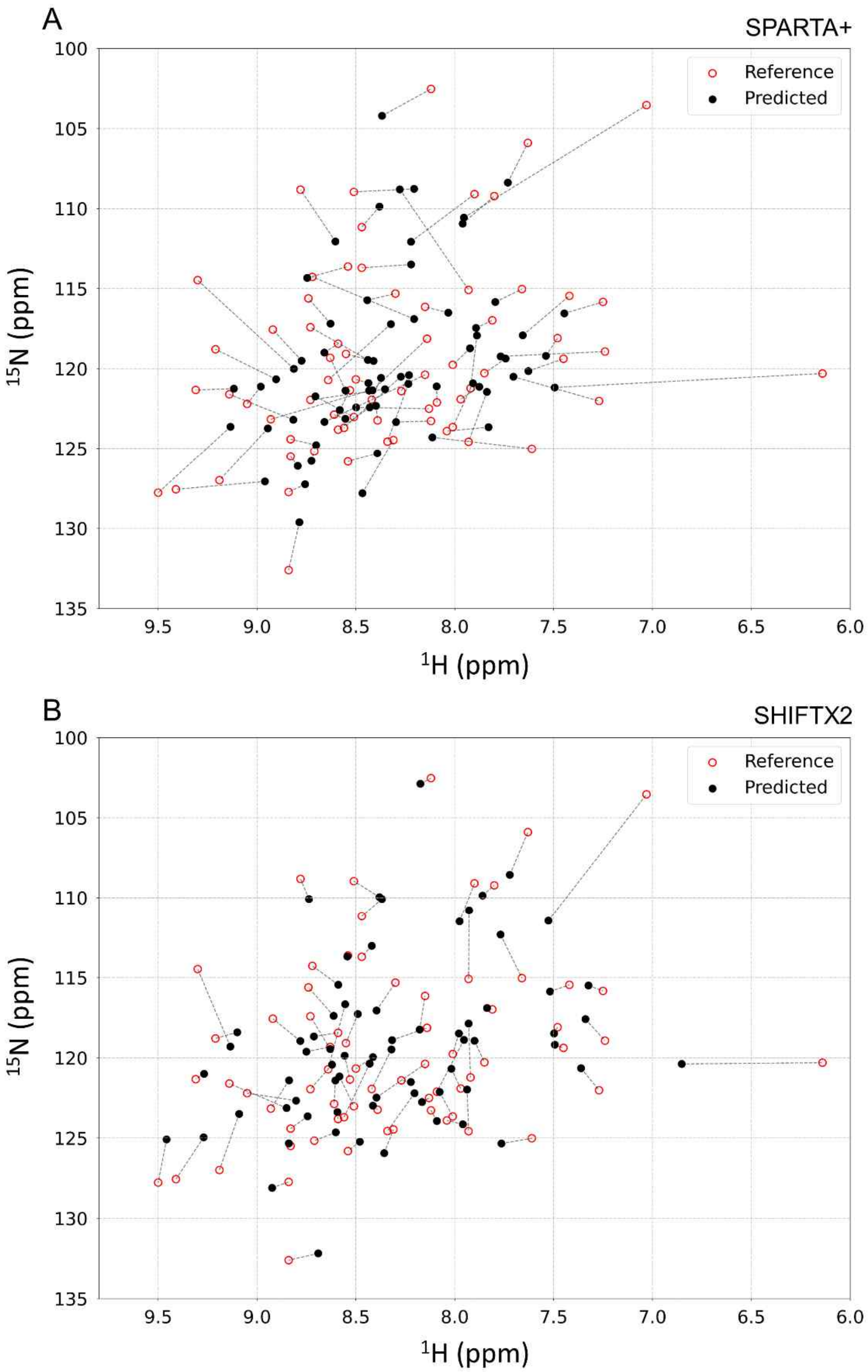

**Figure S14. Comparison between predicted and experimentally-obtained NMR chemical shifts of** chymotrypsin inhibitor 2**.** (A and B) Weighted average of chemical shift prediction using the regression method (black) and chemical shifts obtained using HSQC measurements (red) are shown. NMR chemical shifts were obtained using SPARTA+ (A) and SHIFTX2 (B).

### Supplemental tables

**Table S1.** Average of chemical shift prediction along whole sampled trajectory for Aβ42

| Reference | Solvent Model | UCBSHIFT | | | SPARTA+ | | | SHIFTX2 (10°C) | | |
|---|---|---|---|---|---|---|---|---|---|---|
| | | $Score_{total}$ | $Score_{H}$ | $Score_{N}$ | $Score_{total}$ | $Score_{H}$ | $Score_{N}$ | $Score_{total}$ | $Score_{H}$ | $Score_{N}$ |
| Experiment | Implicit | -.0261 | -.0378 | .6910 | .0306 | .0561 | .5444 | .0116 | .0189 | .6116 |
| | Explicit | -.0605 | -.0750 | .8073 | .0764 | .1094 | .6985 | -.0419 | -.0592 | .7077 |
| BMRB | Implicit | -.2245 | -.3525 | .6367 | | | | | | |
| | Explicit | -.0358 | -.0461 | .7752 | | | | | | |

**Table S2.** Multiple linear regression with regularization process for Aβ42

| Reference | Solvent Model | UCBSHIFT | | | SPARTA+ | | | SHIFTX2 (10°C) | | |
|---|---|---|---|---|---|---|---|---|---|---|
| | | $Score_{total}$ | $Score_{H}$ | $Score_{N}$ | $Score_{total}$ | $Score_{H}$ | $Score_{N}$ | $Score_{total}$ | $Score_{H}$ | $Score_{N}$ |
| Experiment | Implicit | .9456 | .9949 | .9504 | .7942 | .9797 | .8107 | .8745 | .9817 | .8908 |
| | Explicit | .9528 | .9905 | .9619 | .8037 | .9854 | .8156 | .8661 | .9783 | .8853 |
| BMRB | Implicit | .8737 | .9620 | .9082 | | | | | | |
| | | .8968 | .9733 | .9214 | | | | | | |

UCBSHIFT > SHIFTX2 > SPARTA+

**Table S3.** Average of chemical shift prediction along whole sampled trajectory for ubiquitin

| Reference | Solvent Model | UCBSHIFT | | | SPARTA+ | | | SHIFTX2 (10°C) | | |
|---|---|---|---|---|---|---|---|---|---|---|
| | | $Score_{total}$ | $Score_{H}$ | $Score_{N}$ | $Score_{total}$ | $Score_{H}$ | $Score_{N}$ | $Score_{total}$ | $Score_{H}$ | $Score_{N}$ |
| BMRB | Implicit | .7346 | .8040 | .9135 | .2925 | .3783 | .7733 | .5064 | .6229 | .8129 |

**Table S4.** Multiple linear regression with regularization process for ubiquitin

| Reference | Solvent Model | UCBSHIFT | | | SPARTA+ | | | SHIFTX2 (10°C) | | |
|---|---|---|---|---|---|---|---|---|---|---|
| | | $Score_{total}$ | $Score_{H}$ | $Score_{N}$ | $Score_{total}$ | $Score_{H}$ | $Score_{N}$ | $Score_{total}$ | $Score_{H}$ | $Score_{N}$ |
| BMRB | Implicit | .9805 | .9905 | .9899 | .6366 | .7447 | .8548 | .8319 | .9445 | .8808 |

UCBSHIFT > SHIFTX2 > SPARTA+

**Table S5.** Average of chemical shift prediction along whole sampled trajectory for chymotrypsin inhibitor II

| Reference | Solvent Model | UCBSHIFT | | | SPARTA+ | | | SHIFTX2 (10°C) | | |
|---|---|---|---|---|---|---|---|---|---|---|
| | | $Score_{total}$ | $Score_{H}$ | $Score_{N}$ | $Score_{total}$ | $Score_{H}$ | $Score_{N}$ | $Score_{total}$ | $Score_{H}$ | $Score_{N}$ |
| BMRB | Implicit | .0082 | .0859 | .0953 | -.0004 | -.0260 | .0171 | .0420 | .2016 | .2082 |

**Table S6.** Multiple linear regression with regularization process for chymotrypsin inhibitor II

| Reference | Solvent Model | UCBSHIFT | | | SPARTA+ | | | SHIFTX2 (10°C) | | |
|---|---|---|---|---|---|---|---|---|---|---|
| | | $Score_{total}$ | $Score_{H}$ | $Score_{N}$ | $Score_{total}$ | $Score_{H}$ | $Score_{N}$ | $Score_{total}$ | $Score_{H}$ | $Score_{N}$ |
| BMRB | Implicit | .8789 | .9905 | .8873 | .2137 | .6904 | .3095 | .4896 | .9222 | .5309 |

UCBSHIFT > SHIFTX2 > SPARTA+
